# Membrane bending by protein phase separation

**DOI:** 10.1101/2020.05.21.109751

**Authors:** Feng Yuan, Haleh Alimohamadi, Brandon Bakka, Andrea N. Trementozzi, Nicolas L. Fawzi, Padmini Rangamani, Jeanne C. Stachowiak

**Affiliations:** University of Texas at Austin, Department of Biomedical Engineering; University of California San Diego, Department of Mechanical and Aerospace Engineering; Brown University, The Robert J and Nancy D Carney Institute for Brain Science & Department of Molecular Pharmacology, Physiology and Biotechnology; University of Texas at Austin, Institute for Cellular and Molecular Biosciences

## Abstract

Membrane bending is a ubiquitous cellular process that is required for membrane traffic, cell motility, organelle biogenesis, and cell division. Proteins that bind to membranes using specific structural features, such as wedge-like amphipathic helices and crescent-shaped scaffolds, are thought to be the primary drivers of membrane bending. However, many membrane-binding proteins have substantial regions of intrinsic disorder, which lack a stable three-dimensional structure. Interestingly, many of these disordered domains have recently been found to form networks stabilized by weak, multi-valent contacts, leading to assembly of protein liquid phases on membrane surfaces. Here we ask how membrane-associated protein liquids impact membrane curvature. We find that protein phase separation on the surfaces of synthetic and cell-derived membrane vesicles creates a substantial compressive stress in the plane of the membrane. This stress drives the membrane to bend inward, creating protein-lined membrane tubules. A simple mechanical model of this process accurately predicts the experimentally measured relationship between the rigidity of the membrane and the diameter of the membrane tubules. Discovery of this mechanism, which may be relevant to a broad range of cellular protrusions, illustrates that membrane remodeling is not exclusive to structured scaffolds, but can also be driven by the rapidly emerging class of liquid-like protein networks that assemble at membranes.

**Significance Statement:** Cellular membranes take on an elaborate set of highly curved and bent shapes, which are essential to diverse cellular functions from endocytosis to cell division. The prevailing view has been that membrane bending is driven by proteins with curved shapes, which assemble at the membrane surface to form solid scaffolds. In contrast, here we show that proteins which form liquid-like assemblies on membranes are also potent drivers of bending. These “liquid scaffolds” apply compressive stress to the membrane surface, generating a diverse and dynamic family of membrane shapes. These data, which come at a time when liquid-like protein assemblies are being identified throughout the cell, suggest that liquid-like protein assemblies may play an important role in shaping cellular membranes.

## Introduction

From endocytic buds (1) to needle-like filopodial protrusions (2), curved membrane surfaces play critical roles in many cellular processes (3). The energetic cost of creating these highly curved surfaces is considerable, such that spontaneous membrane fluctuations are insufficient to establish and stabilize the shapes of cellular membranes (4). Instead, work during the past two decades has revealed that interactions between proteins and lipids drive membrane curvature (5). Multiple physical mechanisms underlie the ability of proteins to shape membrane surfaces. These include amphipathic helices that insert like wedges into one leaflet of the membrane, creating an inter-leaflet area mismatch that drives curvature (6). Alternatively, proteins with inherently curved membrane binding domains, such as BAR domains, dynamin, and ESCRTs, act as scaffolds that can stabilize curved membrane geometries (7, 8). While each of these mechanisms relies on structured protein domains, we have recently reported that intrinsically disordered proteins, which lack a stable three-dimensional structure, can also be potent drivers of membrane bending (9, 10). Specifically, when non-interacting disordered domains are crowded together in cellular structures, steric repulsion among them drives the membrane to buckle outward, taking on a curved shape.

Interestingly, rather than repelling one another, many disordered proteins have recently been found to assemble together via weak, multi-valent interactions, forming networks that have the physical properties of liquids (11). Notably, recent studies have suggested that liquid-liquid phase separation of membrane-bound proteins plays an important role in diverse cellular processes including nucleation of actin filaments (12), immunological signaling (13), and assembly of virions (14).

How might liquid-liquid phase separation of proteins at membrane surfaces impact membrane curvature? To address this question, we examined phase separation of the N-terminal low complexity domain of fused in sarcoma, FUS LC, on the surfaces of synthetic and cell-derived membrane vesicles. FUS LC was chosen as a model protein for this study because it is among the most thoroughly characterized examples of a domain that undergoes liquid-liquid protein phase separation in solution (15). Here, we assemble FUS LC on membrane surfaces using an N-terminal histidine tag (16) that binds strongly to lipids with Ni-NTA headgroups. As FUS LC accumulated at the membrane surface, we observed protein phase separation in the two-dimensional plane of the membrane, followed by spontaneous inward bending of the membrane, such that protein-lined tubules were created. Similar tubules were observed with two other domains implicated in liquid-liquid phase separation, the low complexity domain of hnRNPA2 (17) and the RGG domain of LAF-1 (18). Interestingly, the tubules had undulating morphologies, similar to a string of pearls. This phenomenon is associated with an area mismatch between the two membrane leaflets (19, 20), suggesting that protein phase separation pulls lipids toward one another, creating a compressive stress on one side of the membrane. In line with this hypothesis, a continuum mechanics model recreated the tubule morphology when a compressive stress was imposed on the outer membrane surface. Further, the model predicted that tubule diameter should increase with increasing membrane rigidity and increasing rigidity ratio, trends confirmed by our experiments. Collectively, these findings suggest that protein phase separation on membrane surfaces generates considerable stresses that can drive the spontaneous assembly of membrane buds and tubules with physiologically relevant dimensions.

## Results

### Protein phase separation on membranes drives assembly of protein-lined tubules

To examine the impact of protein phase separation on membrane surfaces, we combined an N-terminal 6 histidine-tagged version of FUS LC, his-FUS LC, with giant unilamellar vesicles consisting of 93 mol% POPC, 5 mol% Ni-NTA, 2 mol% DP-EG10 biotin for coverslip tethering, and 0.1 mol% Texas Red-DHPE for visualization, Figure 1a. The protein was labeled at the N-terminus with an NHS-reactive dye, Atto 488 for visualization, as described under materials and methods. Samples were imaged using multi-channel, high magnification spinning disc confocal microscopy. When a protein concentration of 0.5 μM was applied to the vesicles, a relatively dim, uniform signal from the protein was observed at the membrane surface, Figure 1b. In contrast, when the protein concentration was increased to 1 μM, more intense regions of fluorescence in the protein channel were observed around the vesicle periphery, Figure 1c. Three dimensional reconstruction of image stacks revealed that these bright regions formed hemispherical domains on the vesicle surfaces, which were surrounded by dimmer regions, Figure 1c, protein panel.

**Figure 1.**
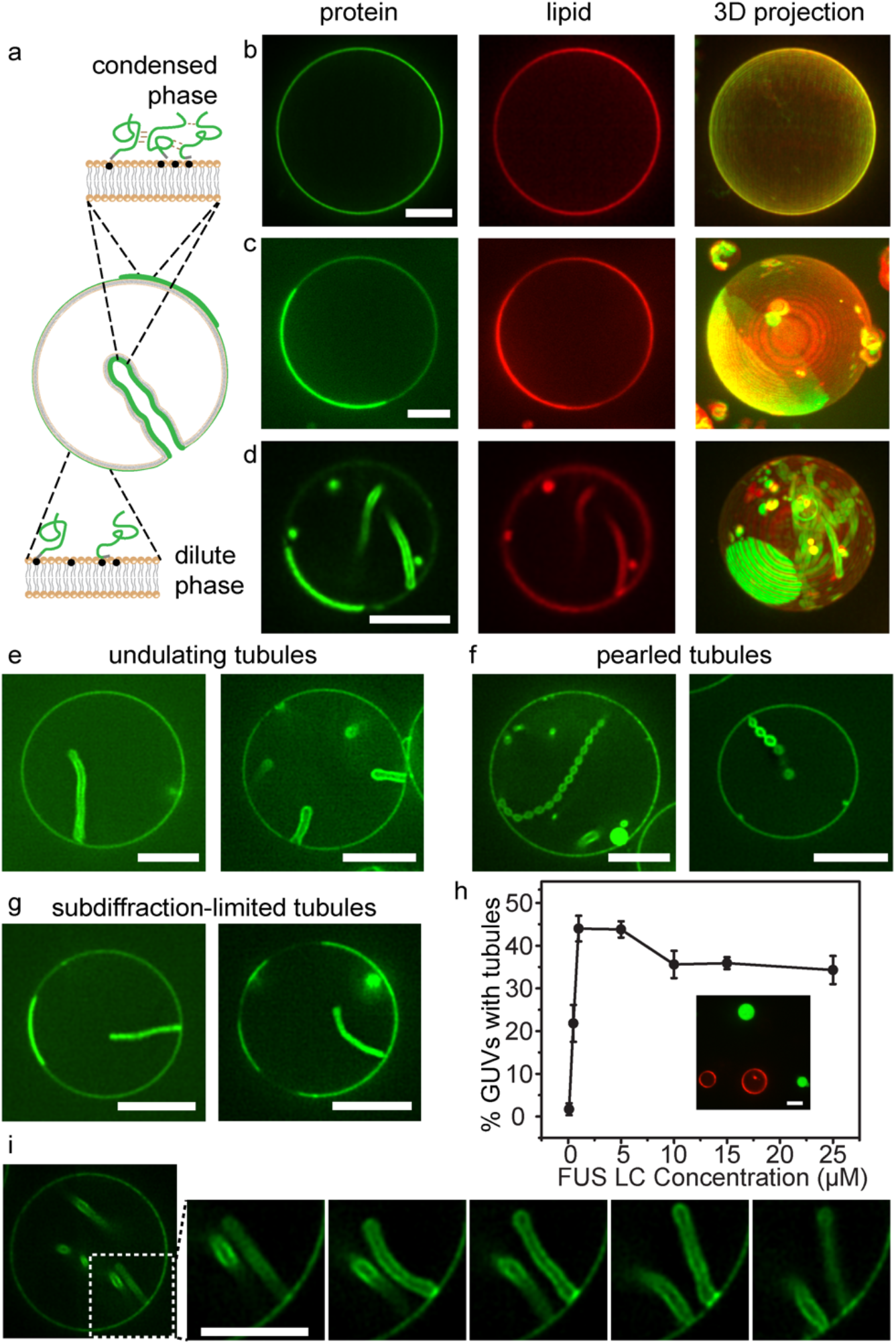
Protein phase separation on membranes drives assembly of protein-lined tubules. (**a**) Pictorial representation of his-FUS LC liquid-liquid protein phase separation on GUV membranes and inward tubule formation. Green lines represent FUS LC proteins. Grey domains indicated 6×histidine tags, and the black dots indicate Ni-NTA lipids. (**b-g)**Representative super-resolution images of GUVs incubated with 0.5 μM (**b**) and 1 μM atto-488 labeled his-FUS LC (**c-g**) in 25 mM HEPES, 150 mM NaCl buffer, pH 7.4. (**b-d**) Representative confocal images (lipid and protein channels) and corresponding maximum intensity projects of GUVs incubated with his-FUS LC. Some GUVs are covered uniformly by the protein (**b**), while others display 2D liquid-liquid phase separation (**c**), which is frequently correlated with the formation of lipid tubules (**d**). (**e-g)**Three kinds of membrane tubule structures were observed: undulating tubules (**e**), tubules consisting of a string of pearls (**f**), and sub-diffraction limited tubules, the structure of which cannot be clearly resolved (**g**). GUV membrane composition: 93 mol% POPC, 5 mol% Ni-NTA, 2 mol% DP-EG10 biotin and 0.1 mol% Texas Red-DHPE. (**h**) The fraction of GUVs displaying inward tubules as a function of his-FUS LC concentration. Data represent mean ± standard deviation; n = 3 independent experiments; N > 100 GUVs were acquired in each replicate. When the addition of his-FUS LC was greater than 5 μM, protein droplets were observed in the surrounding medium (inset in **h**). (**i**) Confocal image series illustrating dynamic fluctuations in tubule shape. All scale bars correspond to 5 μm.

The appearance of these vesicles is remarkably similar to vesicles undergoing phase separation into two coexisting lipid phases (21, 22). In particular, the protein-rich regions in Figure 1c,d have smooth, rounded boundaries, suggesting that they enclose an easily deformable liquid (21). However, the membrane composition used in the present study consisted entirely of unsaturated lipids with melting temperatures well below room temperature, such that phase separation of the underlying lipid membrane was not expected. Further, a control protein that is not involved in protein phase separation, histidine-tagged GFP, covered the surfaces of these vesicles uniformly, Figure S1. These results suggest that the variations in intensity in the his-FUS LC protein channel did not arise from lipid heterogeneity. Instead the FUS LC protein appeared to organize on the two-dimensional membrane surface into protein-rich and protein-poor phases. Notably, the head-labeled lipid probe, Texas Red DHPE, was slightly brighter within the protein-rich regions, likely owing to affinity between the aromatic fluorophore on the lipid headgroup and the FUS LC domain, which is enriched in aromatic tyrosine residues (15). However, photophysical effects of FUS LC on Texas Red could also play a role. To separate the lipid label from FUS LC, we also conducted experiments with a tail group labeled lipid, Texas Red-ceramide. Here enrichment of the labeled lipid in the protein-rich regions was lost, further suggesting that lipid phase separation does not occur in these vesicles, Figure S2.

A few minutes after the addition of his-FUS LC, we observed that many of the vesicles developed lipid tubules spontaneously. These tubules originated at the surfaces of the vesicles and protruded into the vesicle lumen, such that they were lined by the his-FUS LC protein, Figure 1d. Some of the tubules had an undulating, wavy appearance, Figure 1d, e, while others formed a series of tight spheres, resembling a string of pearls, Figure 1f. Still others were so slender that their morphology could not be precisely determined, Figure 1g. In some instances, tubules remain associated with protein-rich membrane domains, Figure 1g, while in other cases, the domains appear to have been consumed, transforming fully into tubules, Figure 1f.

Tubules were observed more frequently as the concentration of his-FUS LC increased, Figure 1h, Table S1. Specifically, less than 2% of vesicles formed lipid tubules in the presence of 0.1 μM FUS LC, while 22% and 44% formed tubules after addition of 0.5 μM and 1 μM FUS LC, respectively. However, for protein concentrations above 1 μM, the fraction of vesicles with tubules reached a plateau, likely owing to the appearance of three-dimensional protein-droplets in the surrounding solution, which did not appear to be membrane associated, Figure 1h (inset). These droplets likely compete with the membrane surface for protein molecules, limiting the further accumulation of protein on the membrane surface.

Importantly, dynamic fluctuations were observed in the morphology of the tubules, suggesting that the protein layer on the membrane surface remained highly deformable, rather than assembling into a rigid scaffold, Figure 1i, Supplementary Movie 1. Additionally, domains of the protein-depleted phase had rapidly fluctuating boundaries and diffused randomly within the protein-enriched phase, observations which further demonstrate the fluid-like nature of the protein-rich phase, Supplementary Movie 2. To further quantify the relationship between protein concentration and tubule formation, we next varied the strength of protein-protein and protein-membrane interactions and observed the impact on the membrane tubules.

### Assembly of lipid tubules depends on the strength of protein-protein and protein-membrane interactions

The membrane tubules in Figure 1 appear to emerge from the protein-rich regions of the membrane surface, suggesting that they rely on self-association of membrane-bound his-FUS LC molecules. We would expect that the ability of these proteins to come together on membrane surfaces depends on both the extent of protein-membrane binding and the strength of protein-protein interactions. Therefore, the assembly of membrane tubules likely depends upon these parameters. To vary the extent of protein-membrane binding, we varied the concentration of Ni-NTA-DOGS lipids in the membrane vesicles. To vary the strength of interactions between his-FUS LC proteins, we varied the concentration of sodium chloride in the solution. This approach is based on published studies showing that the saturation concentration for liquid-liquid phase separation of FUS LC decreases as sodium chloride concentration increases, resulting in enhanced phase separation (16).

Holding the concentration of his-FUS LC constant at 1 μM, we mapped the prevalence of two-dimensional protein phase separation and lipid tubules as a function of both NaCl concentration (50 mM – 250 mM) and the concentration of Ni-NTA-DOGS lipids (2-15 mol%), Figure 2a-d. We observed that increasing either parameter led to an increase in both the fraction of vesicles exhibiting phase separation, Figure 2e and Table S2, and the fraction of vesicles exhibiting lipid tubules, Figure 2f and Table S3. Plotting the fraction of phase separated vesicles versus the fraction of vesicles with lipid tubules, reveals a sharp transition to strong tubule formation when approximately 25% or more of the vesicles display phase separation, Figure 2g, Pearson’s correlation coefficient, 0.8.

**Figure 2.**
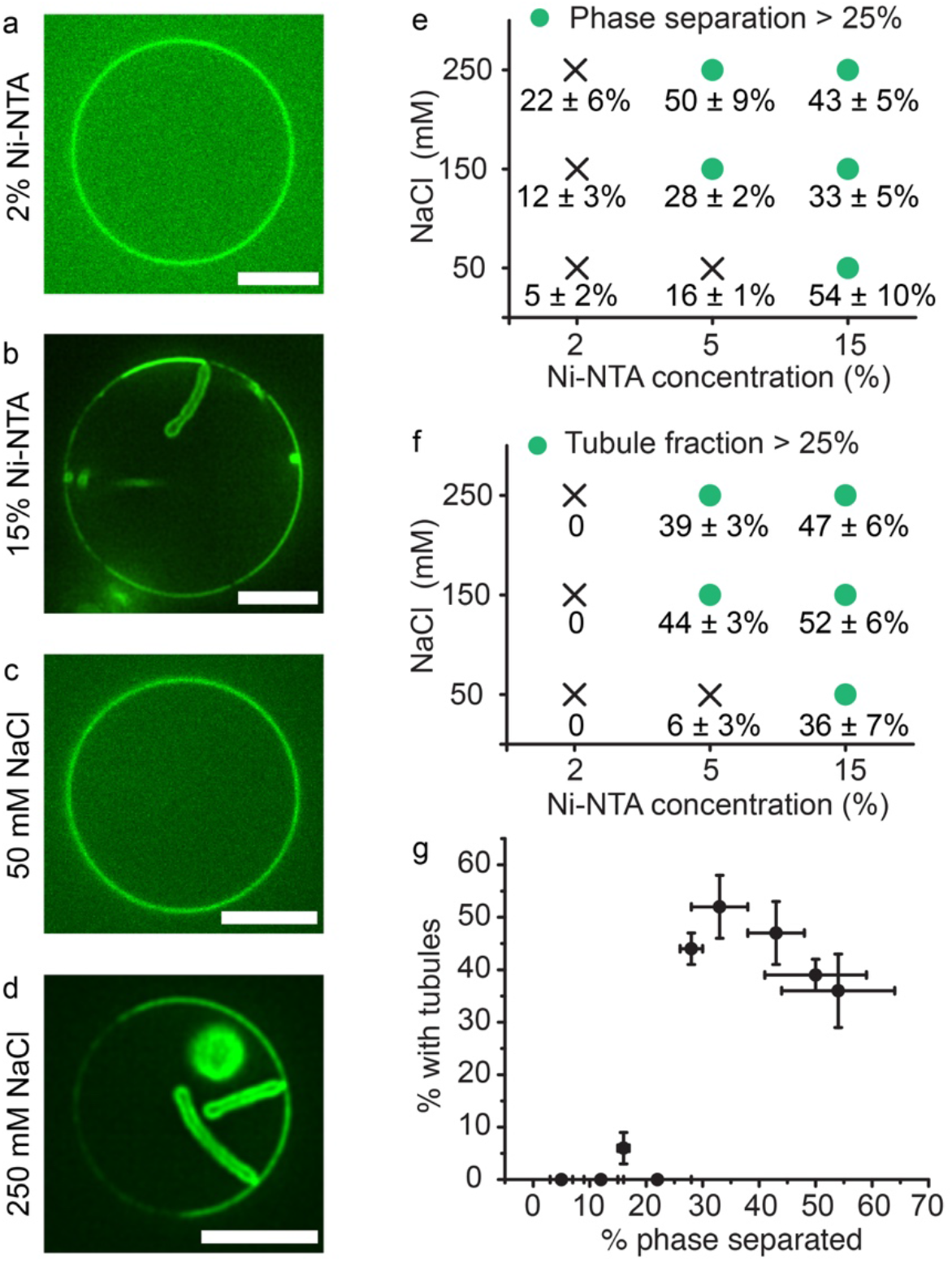
Protein phase separation and tubule formation depend upon the concentration of membrane-bound proteins and the strength of protein-protein interactions. (**a**) Representative confocal images of FUS LC bound to GUVs (composition: 96 mol% POPC, 2 mol% Ni-NTA, 2 mol% DP-EG10-biotin, 0.1 mol% Texas Red-DHPE) containing 2 mol% Ni-NTA, and (**b**) GUVs (composition: 83 mol% POPC, 15 mol% Ni-NTA, 2 mol% DP-EG10-biotin, 0.1 mol% Texas Red-DHPE) containing 15% Ni-NTA. GUVs were incubated with 1μM Atto-488 labeled his-FUS LC in 25 mM HEPES, 150 mM NaCl pH 7.4 buffer. (**c, d**) Representative images of GUVs (93 mol% POPC, 5 mol% Ni-NTA, 2 mol% DP-EG10 biotin and 0.1 mol% Texas Red-DHPE made in 560 mOsmo glucose solution) incubated with 1μM atto-488 labeled FUS LC in 25 mM HEPES pH 7.4 buffer containing (**c**) 50 mM and (**d**) 250 mM NaCl, respectively. Glucose was added to the buffers accordingly to maintain osmotic pressure balance. All scale bars represent 5 μm. (**e, f**) Percentage of GUVs displaying (**e**) inward lipid tubules and (**f**) protein phase separation as a function of Ni-NTA content, and NaCl concentration. Green dots indicate fractions exceeding 25%. (**g**) Percentage of GUVs with inward tubules as a function of percentage of GUVs with phase separation. Here the Pearson’s correlation coefficient between phase separation and tubule formation was 0.8. Data are shown as mean value ± standard deviation. N > 100 GUVs were analyzed cumulatively from three independent replicates for each condition.

These results demonstrate that formation of protein-lined lipid tubules is strongly correlated with phase separation of his-FUS LC on membrane surfaces. However, it remains unclear why phase separation on membrane surfaces drives the membrane to bend inward, toward the lumen of the vesicle. In order to understand this phenomenon, we developed a continuum mechanical model of membrane bending in the presence of protein phase separation.

### A continuum mechanics model predicts tubule shape and dependence of tubule diameter on membrane bending rigidity

The morphologies of the tubules that we have observed can provide insights into the mechanism by which protein phase separation drives membrane bending. Some tubules consist of a well-defined “string of pearls” in which spherical shapes are separated by thin necks, Figures 1f. Other tubules have an undulating morphology in which the “pearls” are less well defined, with some tubules being nearly cylindrical, Figure 1e, 2b, d. This set of shapes - pearls, undulations, and cylinders - can be classified as Delaunay surfaces (23), which have a constant, non-zero mean curvature, Figure 3a. Unduloids are surfaces of revolution of an elliptic catenary (23, 24). With small changes in geometric parameters, a range of unduloid surfaces can be constructed (23), Figure 3a. More importantly, Delaunay surfaces, particularly unduloids and their variants, are known to minimize the Helfrich energy for membrane bending (24). The radius and shape of the unduloids depends on a single dimensionless parameter 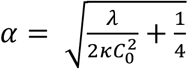, where λ is the membrane tension, *κ* is the bending modulus, and *C*_0_ is the spontaneous curvature. When *α* = 0.75, the membrane resembles a cylinder and for *α* > 0.75, the membrane resembles an unduloid, Figure 3a.

**Figure 3.**
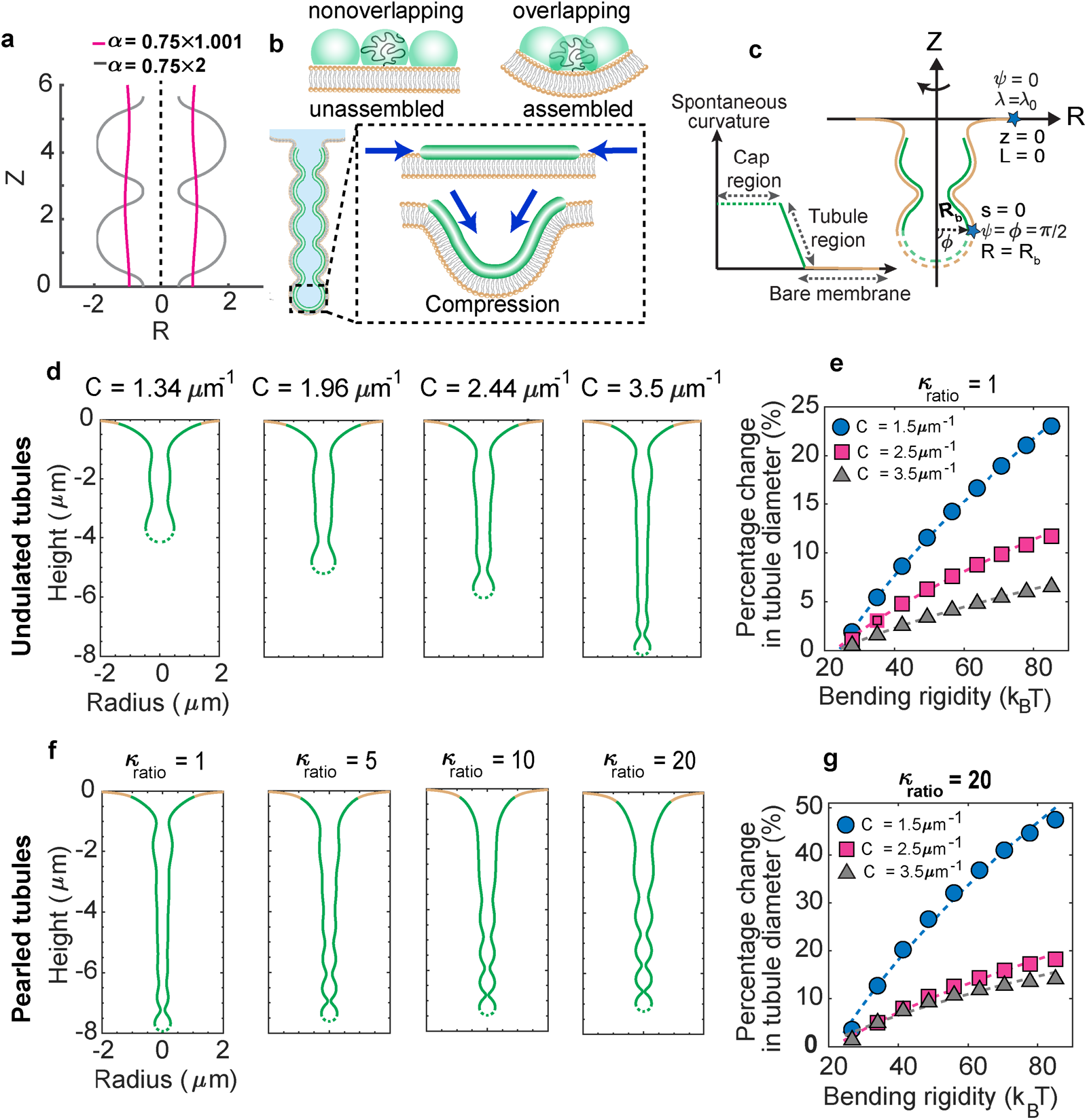
Mechanical model of undulating and pearled tubule formation. (**a**) Unduloid-like shapes solution for Helfrich energy minimization at different values of non-dimensional parameter, *α*. For α~0.75, the membrane takes on a cylindrical shape (purple line); for α>0.75, the unduloid becomes a sphere similar to a string of pearls (gray line). (**b**) Schematic depiction of membrane tubule formation due to the compressive stresses applied by liquid-liquid phase separation on the membrane. (**c**) Schematic of the axisymmetric simulations depicting the simulation domain and the boundary conditions. The yellow region represents the bare membrane and the green region is the area coated by the proteins. The dashed lines indicate the cap of the tubule, assumed to have a constant curvature. The inset shows the spontaneous curvature distribution along the tubule region used to model the membrane shape. (**d**) Undulating tubules minimize the membrane bending energy as the spontaneous curvature increases for uniform bending rigidity of the membrane (*κ* = 80 k_B_T). (**e**) Percentage of change in the tubule diameter ((D-D*_κ_* _= 25 kBT_)/D*_κ_* _= 25 kBT_) as a function of the bending rigidity for three different values of spontaneous curvature. The dashed lines show a square root dependence on the bending modulus by fitting to the curve 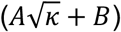 where for the gray line A=5.4, B=−26.44; for the pink line A= 2.71, B=−12.9; and for the blue line, A=1.53 and B=−7.4. (**f**) Pearled tubules minimize the bending energy of the membrane for heterogeneous membrane rigidity (*κ_ratio_*= *κ_protein-domain_*/*κ_bare membrane_*), C_0_ = 3.5 *μ*m^-1^. (**g**) Percentage of change in the tubule diameter ((D-D*_κ_* _= 25 kBT_)/D*_κ_* _= 25 kBT_) as a function of the bending rigidity for three different values of spontaneous curvature for *κ_ratio_*= 20. The dashed lines are the fitted curve 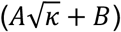 where for the gray line; A=10.98, B=−51.31, for the pink line; A= 4.22, B=−19.58, and for the blue line; A=3.1 and B=−13.

Tubules with unduloid-like morphologies are known to arise when there is an area mismatch between the inner and outer leaflets of the lipid bilayer, such that the membrane has a finite spontaneous curvature (25). For example, addition of lipids (19), polymers (26), and proteins (27) to the surfaces of membrane vesicles have each been shown to produce such tubules. However, in these cases, the tubules protruded outward from the membrane surfaces, as would be expected when the area of the outer leaflet exceeds that of the inner leaflet. In contrast, we observe tubules that protrude inward from the membrane surface, suggesting that protein phase separation reduces the area of the outer leaflet relative to that of the inner leaflet, Figure 3b. We might expect such a reduction in area if phase separation of his-FUS LC peptide generates compressive forces at the membrane surface. Specifically, the conformational freedom of any tethered peptide chain should increase with increasing distance from the membrane surface (28). As a result, protein residues near the free end of the chain are more likely to interact with one another than are residues near the tethered end of the chain. Inward bending of the membrane would be expected to increase overlap between the parts of the peptide chains that are distant from the membrane surface, Figure 3b. In this way, protein phase separation would be expected to stabilize inward protrusions of the membrane.

To examine these ideas, we developed a model in which a compressive stress was applied to one leaflet of a lipid bilayer. The membrane bending energy was modeled using the Helfrich framework (29). The area difference between the two leaflets was modeled using a locally specified spontaneous curvature for simplicity in simulations, Figure 3c. See the supplemental information for detailed model assumptions and derivations. The spontaneous curvature effectively represents the stresses due to the area difference between the two leaflets (30). The governing equations were solved in an axisymmetric parametrization for ease of computation to demonstrate the principles underlying the formation of undulating and pearled tubules.

We first simulated a domain of fixed area and homogeneous bending rigidity that included the protein enriched phase and the surrounding protein depleted phase. Our results showed that increasing the spontaneous curvature in the protein-rich phase resulted in the formation of undulating tubules, Figure 3d, similar to those observed in experiments, Figure 1. Furthermore, the simulations predicted that the tubule diameter would increase linearly in proportion to the square root of the bending modulus, Figure 3e. The bending energy corresponding to the formation of the undulating and pearled tubules is shown in Figure S3.

It is likely that the protein enriched phase has an increased bending rigidity compared to the protein depleted phase, owing to the higher density of protein contacts. Therefore, we next asked if the ratio of bending rigidities in the attached protein layer and the underlying membrane layer could impact the tube diameter. We defined the ratio of bending rigidities, 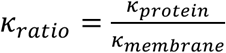 and varied the ratio in the range of 1-20, where *κ_ratio_* = 1 denotes uniform bending rigidity. With increasing *κ_ratio_*, we observed that the tubules took on a more clearly defined pearled morphology, Figure 3f,g and Figure S4, similar to those observed in some of our experiments, Figure 1f. We next sought to test these predictions.

### Tubule diameter varies with membrane bending rigidity and salt concentration

The continuum model predicted that the radii of the tubules should increase in proportion to the square root of the membrane bending rigidity. To examine this prediction, we measured the diameters of the resolvable lipid tubules formed by assembly of his-FUS LC on membrane surfaces, as a function of membrane bending rigidity, Figure 4a-e, Table S4. As membrane bending rigidity was increased from approximately 20 k_B_T to approximately 76 k_B_T, through variations in membrane lipid composition, we observed a substantial increase in membrane tubule diameter from 240 ± 100 nm (S. D.) to 400 ± 190 nm (S. D.), Figure 4e. For each lipid composition, tubules with both pearled and undulating morphologies were observed, Figure 4a-d. Further, the data were reasonably well fit by a curve in which tubule diameter was proportional to the square root of bending rigidity, in agreement with the predictions of the simulation, compare Figure 4f, Figure 3e, g. Here optical reassignment during spinning disc confocal microscopy, followed by deconvolution was used to increase the optical resolution to better than 150 nm (31).

**Figure 4.**
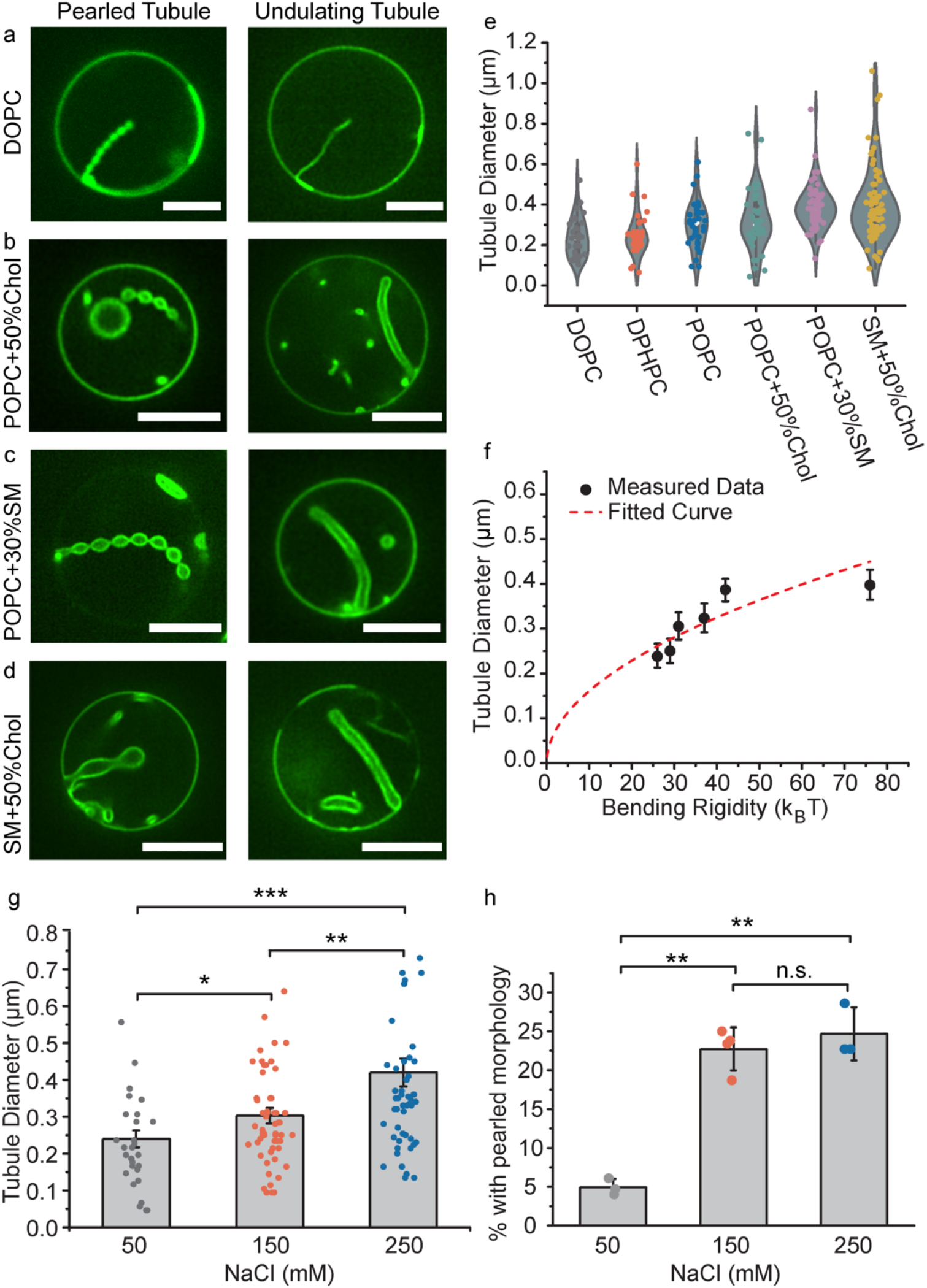
Tubule diameter varies with membrane bending rigidity and salt concentration. (**a-f**) Six groups of GUVs with different compositions (listed in Table S4 in Materials and Methods) were used to vary membrane bending rigidity. GUVs were incubated with 1μM atto-488 labeled his-FUS LC in 25 mM HEPES, 150 mM NaCl pH 7.4 buffer. (**a-d**) Representative super-resolution confocal images tubules with pearled (left panel) and undulating morphologies (right panel), from GUVs consisting primarily of (**a**) DOPC, (**b**) POPC + 50% Chol, (**c**) POPC + 30% SM, and (**d**) SM + 50% Chol. All scale bars are 5 μm. (**e)**Violin plot showing the measured tubule diameter distribution for tubules formed using each GUV composition. (**f)**GUV tubule diameter as a function of membrane bending rigidity. Data points from left to right represent DOPC, DPHPC, POPC, POPC + 50% Chol, POPC + 30% SM, and SM + 50% Chol, respectively. Data are displayed as mean ± standard error from at least 60 tubules per composition, gathered during 3 independent experiments. The measured tubule diameters increase roughly as the square root of membrane bending rigidity (red dash line, R^2^ = 0.64). (**g**) Bar chart displaying average tubule diameter under different NaCl concentrations. GUVs (composition: 83 mol% POPC, 15 mol% Ni-NTA, 2 mol% DP-EG10-biotin and 0.1% Texas Red-DHPE) were incubated with 1μM atto-488 labeled his-FUS LC in 25 mM HEPES, pH 7.4 buffer with corresponding NaCl concentration under iso-osmotic conditions. Error bars correspond to standard error. Each point is a mean value of diameters measured at three positions along the same tubule. N > 100 GUVs were acquired cumulatively from three independent replicates for each condition. (**h**) Fraction of tubules that displayed a pearled morphology as a function of NaCl concentration Data are displayed as mean ± standard deviation from three independent experiments (n = 3) on separate preparations of vesicles, with cumulatively N > 100 vesicles categorized. Brackets show statistically significant comparisons using an unpaired, 2-tailed student’s t test. * represents p < 0.05, ** represents p < 0.01, *** represents p < 0.001, and n.s. indicates a difference that was not statistically significant.

A second prediction from our simulation is that the tubule diameter should increase as the rigidity of the protein-rich phase increases, while the rigidity of the underlying membrane is held constant. To test this prediction, we examined the impact of sodium chloride concentration on tubule diameter. Increasing sodium chloride concentration has been previously shown to increase the strength of interactions between FUS LC molecules in condensed phases (15). Therefore, we inferred that his-FUS LC might assemble into a more rigid protein layer at high salt concentration.

As the sodium chloride concentration increased from 50 mM to 250 mM, we observed an increase in tubule diameter of approximately 75%, from 240 ± 120 nm to 420 ± 280 nm (S. D.), in qualitative agreement with simulation results, compare Figure 4g with Figure 3g. Further, the incidence of tightly pearled tubules increased significantly as NaCl concentration increased from 50 mM to 250 mM, in agreement with simulations, Figure 4h. A constant osmotic pressure was maintained in all experiments, see methods. Collectively these data suggest that protein phase separation applies a compressive stress to the membrane surface, resulting in assembly of protein tubules directed inward from the membrane surface.

### Membrane bending by protein phase separation is a general phenomenon that can be driven by diverse protein domains

The model we have developed does not take into account the specific amino acid sequence of the FUS LC domain or the particular types of molecular interactions that drive the protein to phase separate. Instead we have described tubule formation as a general process that could arise whenever protein phase separation occurs at the membrane surface. Therefore, we next asked whether the ability to drive lipid tubule formation is specific to FUS LC, or whether it is a general property of membrane-bound domains that undergo liquid-liquid phase separation. To address this question, we evaluated two additional domains known to be involved in liquid-liquid phase separation, the low complexity domain of hnRNPA2 (hnRNPA2 LC), a protein involved in RNA processing and transport granule formation (17), and the RGG domain of LAF-1 (LAF-1 RGG), a DDX3 RNA helicase found in *C elegans* P granules (18). Both proteins contained N-terminal histidine tags, which we used to bring them to the membrane surface, as we did with FUS LC, Figure 5.

**Figure 5.**
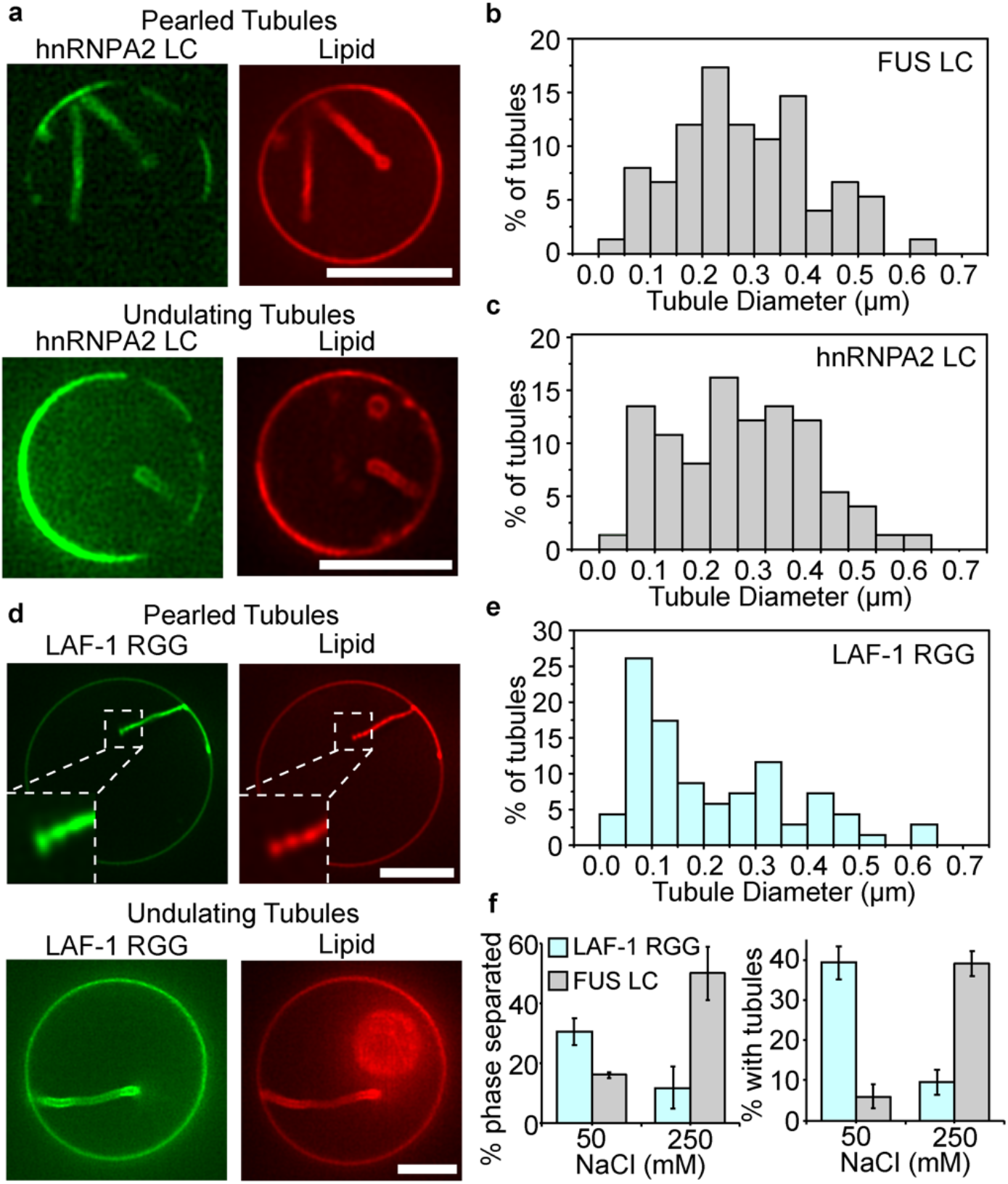
hnRNPA2 LC and Laf-1 RGG domains drive formation of inwardly directed membrane tubules with similar morphologies to those formed by FUS LC. (**a**) his-hnRNPA2 LC at a concentration of 1 μM drove formation of inwardly directed tubules with pearled and undulating morphologies when introduced to GUVs consisting of 83 mol% POPC, 15 mol% Ni-NTA, 2 mol% DP-EG10 biotin and 0.1 mol% Texas Red-DHPE. (**b**) Distribution of tubule diameters formed upon exposure of GUVs to his-FUS LC, 75 total tubules. (**c**) Distribution of tubule diameters form upon exposure of GUVs to his-hnRNPA2 LC, 75 total tubules. (**d**) his-Laf-1 RGG at a concentration of 1 μM drove formation of inwardly directed tubules with pearled and undulating morphologies when introduced to GUVs of the same composition as in a. (**e**) Distribution of tubule diameters formed upon exposure of GUVs to his-Laf-1 RGG, 70 total tubules. (**f**) The fraction of vesicles exhibiting two-dimensional protein phase separation and tubule formation by his-Laf-1 RGG decreased with increasing salt concentration. This is the opposite trend of that observed for vesicles exposed to his-FUS LC, data repeated from Figure 2e, f, for comparison. Error bars represent standard deviation of 3 trials, with cumulatively N > 300 GUVs analyzed. The scale bar in a, d is 5 μm.

Similar to FUS LC, hnRNPA2 LC is a prion-like domain composed primarily of polar and aromatic residues, which contains relatively few aliphatic residues and is depleted in charged residues (17). Both FUS LC and hnRNPA2 LC have an increased propensity to undergo liquid-liquid phase separation as the ionic strength of the surrounding medium increases (15, 17). Based on these similarities, we might expect hnRNAPA2 LC and FUS LC to have similar interactions at the membrane surface and therefore to behave similarly in our assays. As expected, when hnRNPA2 LC was added to giant vesicles at a concentration of 1 μM, inwardly directed lipid tubules with undulating and pearled morphologies were observed, Figure 5a. Further, the distribution of tubule diameters was similar between hnRNPA2 LC and FUS LC, Figure 5b,c.

In contrast to hnRNPA2 LC, LAF-1 RGG has a quite different sequence composition when compared to FUS LC. In particular, LAF-1 RGG is dense in charged residues such as arginine and aspartic acid (18). In this way, increasing the ionic strength of the surrounding medium opposes liquid-liquid phase separation of LAF-1 RGG (18), suggesting that the dominant driving force for liquid-liquid phase separation is electrostatic attraction between oppositely charged residues. To examine the impact of these differences on the formation of membrane tubules, we added 1 μM of LAF-1 RGG to giant vesicles. Interestingly, we observed inwardly directed lipid tubules, which were similar to those formed by FUS LC and hnRNPA2 LC, Figure 5d. The diameters of tubules formed by the three proteins covered approximately the same range, though tubules formed by LAF-1 RGG had a somewhat smaller average diameter, Figure 5e. Importantly, the fraction of giant vesicles that displayed lipid tubules upon exposure to LAF-1 RGG decreased with increasing salt concentration. This trend, which is the opposite of what we observed for FUS LC (Figure 5f), is expected owing to the ability of high ionic strength solutions to screen the electrostatic interactions that support liquid-liquid phase separation of LAF-1 RGG (18).

Collectively, these results demonstrate that the ability of liquid-liquid phase separation at membrane surfaces to drive inward membrane protrusions is a general phenomenon that is not dependent on the specific molecular interactions that drive each protein to phase separate. Instead, liquid-liquid phase separation itself, rather than a particular pattern of electrostatic or hydrophobic interactions between proteins and lipids, appears to be responsible for generating the compressive stress that drives membrane deformation.

### Protein phase separation drives tubule formation from cell-derived membranes

We next asked whether protein phase separation at membrane surfaces is sufficient to drive remodeling of cellular membranes. To address this question, we derived membrane vesicles from the plasma membranes of mammalian retinal pigmented epithelial (RPE) cells. To facilitate binding of FUS LC to the surfaces of these vesicles, we engineered the donor cells to express a chimeric transmembrane protein that consisted of the transmembrane domain of the transferrin receptor, fused to an extracellular blue fluorescing protein, BFP, domain for visualization. This chimera displayed a nanobody against GFP on the cell surface, Figure 6a,b. Membrane blebs extracted from these cells also displayed the nanobody on their surfaces, which facilitated the recruitment of GFP-tagged proteins, Figure 6a,b. Adding soluble GFP domains to the solution surrounding these blebs resulted in GFP being strongly concentrated at the bleb surfaces, Figure 6c. Notably, the GFP signal appeared to separate into brighter and dimmer regions on the surfaces of some of the blebs. This separation within blebs has been observed previously (32), and is thought to arise from lipid phase separation, in which the transferrin receptor transmembrane domain is known to prefer the liquid disordered membrane phase (33).

**Figure 6.**
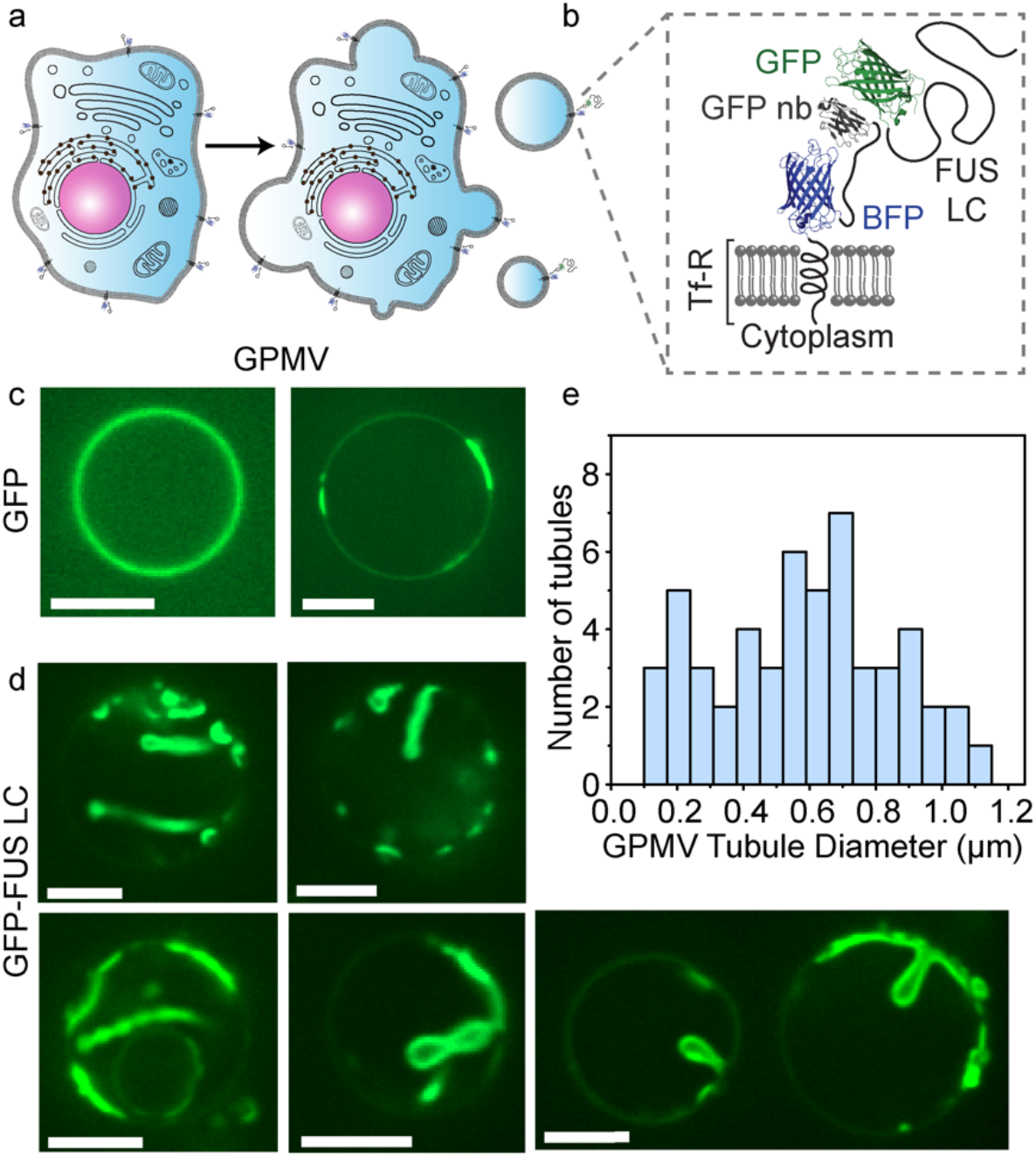
Protein phase separation can drive tubule formation from cell-derived membranes. (**a**) Cartoon showing extraction of giant plasma membrane vesicles (GPMVs) from donor RPE cells. (**b**) Schematic of the architecture of the membrane receptor and ligand protein. GFP-FUS LC is recruited to the GPMV membrane by binding to a GFP nanobody displayed on the cell surface. (**c**) Confocal images of GPMVs incubated with 2 μM GFP and (**d**) GFP-FUS LC in buffer containing 10 mM HEPES, 2 mM CaCl_2_, 150 mM NaCl, pH 7.4. Scale bar = 5 μm. (**e**) Distribution of diameters of tubules formed from GPMVs. N= 50 tubules measured.

When a GFP-tagged version of FUS LC, GFP-FUS LC (34), was introduced to blebs taken from the same donor cells, the GFP signal was similarly concentrated at the bleb surfaces, Figure 6d. However, the surfaces of blebs exposed to GFP-FUS LC did not remain flat. Instead, regions of the bleb surfaces with intense GFP signal bent inward, creating protein-lined membrane buds and tubules. Many of the tubules had pearl-like and undulating morphologies, similar to tubules formed by exposure of synthetic vesicles to his-FUS LC, compare Figures 1 and 6d. The diameter of the tubules ranged broadly from 150 nm to more than 1 μm, Figure 6e. Here, the average tubule diameter, 570 ± 260 nm (S. D.), was somewhat larger than that of tubules formed from synthetic membranes. This difference could arise from the enhanced bending rigidity of cell-derived membranes, which contain a high density of transmembrane proteins. Alternatively, the GFP-FUS LC protein, which has been observed to form gel-like assemblies in solution (34, 35), may increase the rigidity of the protein layer. Nonetheless, the range of curvatures observed in these cell-derived vesicles encompasses that of many cellular structures including filopodia, dendritic spines, phagosomes, and many organelles (3). These results demonstrate that liquid-liquid phase separation of membrane-bound proteins is sufficient to deform complex, cell-derived membranes. Additionally, because these experiments use an antibody-antigen interaction to bring FUS LC to the membrane surface, rather than a histidine tag, these results show that histidine-lipid interactions are not required for membrane bending by liquid-liquid phase separation.

## Discussion

Here, we demonstrate that protein phase separation at membrane surfaces can drive the assembly of protein-lined membrane tubules of physiologically relevant dimensions. This mechanism is physically distinct from membrane bending by solid scaffolds, which include the rigid, tubular assemblies of BAR domains, dynamin, and shiga toxin, as well as the cage-like geometries of protein coats formed by clathrin, COPII, and many viral capsids (7, 36–38). In contrast, we show that a family of model proteins that form liquid-like assemblies can drive the formation of membrane tubules with dynamic cylindrical and unduloid morphologies, Figure 1, Supplementary Movie 1. These results illustrate that increasing the spontaneous curvature of a membrane, which is the fundamental requirement for membrane bending (4, 39–42), is not exclusive to structured scaffolds, but can also arise from liquid-like protein interactions that generate stresses at membrane surfaces. Using the liquid scaffolding mechanism, cytosolic proteins that phase separate at membrane surfaces could contribute to outward membrane protrusions such as filopodia, dendritic spines, viral buds, and cilia. In contrast, proteins and receptors that assemble into liquid scaffolds on the outer cell surface could contribute to structures that bud into the cell, such as endocytic vesicles.

The inward tubule formation observed here in response to liquid-liquid phase separation is in direct contrast to the outwardly protruding tubules generated by repulsive interactions among self-avoiding disordered domains found in endocytic proteins (9, 10). These two sets of observations can be understood as two extremes of the same mechanism. Specifically, the membrane protein composite can be thought as two layers of a 2D fluid, one layer consisting of lipids and the other consisting of proteins. Many studies have shown that lipid bilayers can only be stretched or compressed by a few percent (43, 44). In contrast, the protein layer is capable of dramatic changes in density. When self-avoiding domains become crowded on the membrane surface, they push each other apart. As the protein layer expands, the nearly inextensible lipid bilayer is forced to bend outward. In contrast, when self-interacting proteins undergo liquid-liquid phase separation on the membrane surface, the protein layer contracts, forcing the nearly incompressible lipid bilayer to bend inward. Similar behavior has been observed in simplified models of biological tissues such as intestine (45) and brain (46), where tissues fold owing to the differential compressibility of adjacent two-dimensional layers (47), suggesting a common mechanism in soft matter. While structured protein scaffolds are known to induce anisotropic spontaneous curvature, liquid-like scaffolds arising from assembly of disordered proteins are likely to induce isotropic spontaneous curvature. Notably, the formation of tubules and pearls due to anisotropic protein curvatures have been studied extensively using mechanical models (48–51).

What advantage might a liquid scaffold offer for membrane remodeling? We speculate that the lower energy barriers to assembly and disassembly associated with a liquid may allow the membrane greater freedom to deform into a variety of shapes and dimensions, rather than the more narrowly defined set of geometries observed for most structured scaffolds. Indeed many curved membrane structures, from cytoskeletal protrusions (52) to the endoplasmic reticulum (53), are known to have heterogenous and dynamic morphologies. In particular, the unduloid morphology reported here has been observed in the endosomal networks of plants (54). In light of the ongoing discovery of liquid-like behavior in many membrane-bound protein networks (55), the ability of protein phase separation to shape membranes has the potential to impact membrane-associated processes throughout the cell.

## Acknowledgements

This research was supported by the National Institutes of Health through grants R01GM112065 (J.C.S.), R01GM132106 (P.R.), R01GM118530 (N.L.F), R01NS116176 (N.L.F.), NSF 1845734 (N.L.F.), and by the Human Frontiers Science Program RGP0045/2018 (N.L.F.). The plasmid DNA for GFP FUS LC was provided by the lab of S. McKnight, UTSW.

