## Supplementary Material for "Membrane bending by protein phase separation"

### Supplementary Figures, Tables, and Movie Captions

Please note: Supplementary Figures S3-S4 are integrated with the model description at the end of this document.

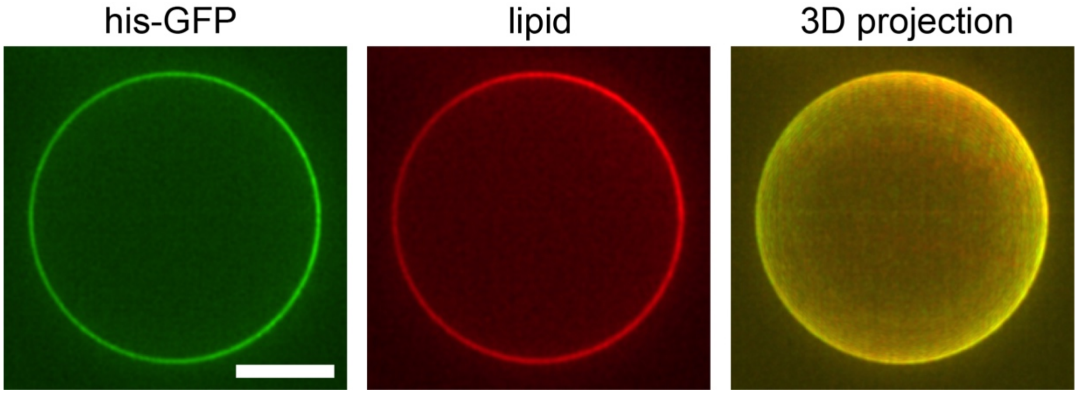


**Figure S1**. Representative super-resolution images of GUVs incubated with 1$\mu$M of his-GFP in 25 mM HEPES, 150 mM NaCl pH 7.4 buffer. GUV consists of 93 mol% POPC, 5 mol% Ni-NTA, 2 mol% DP-EG10-biotin and 0.1 mol% Texas Red-DHPE. Scale bar = 5 $\mu$m.

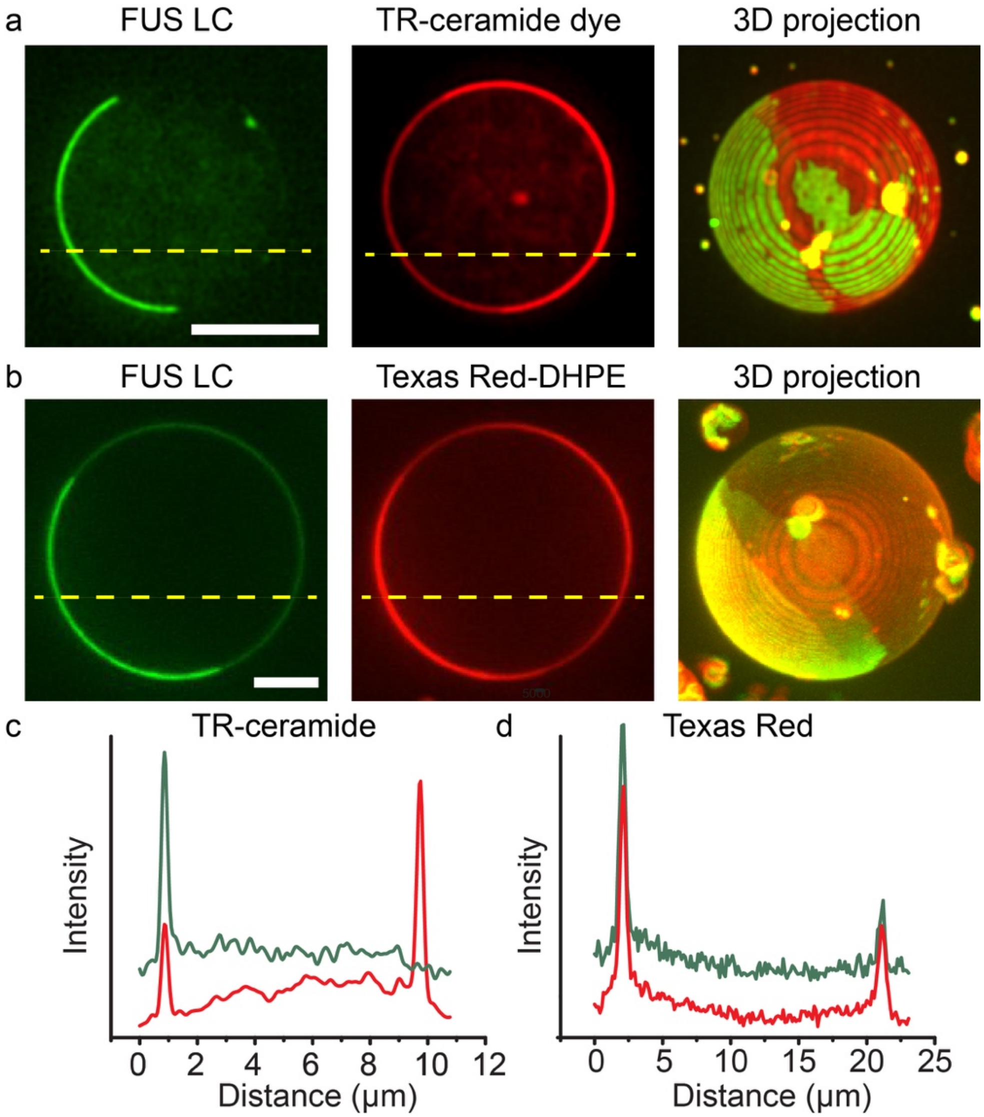


**Figure S2**. Less dye lipids enriched in protein phase separation region when labeled with TR-ceramide. **a, b** Representative super-resolution images of GUVs labeled with TR-ceramide (**a**) and Texas Red-DHPE (**b**) incubated with 1$\mu$M of his-FUS LC in 25 mM HEPES, 150 mM NaCl pH 7.4 buffer. GUVs in panel **a** consist of 83 mol% POPC, 15 mol% Ni-NTA, 2 mol% DP-EG10-biotin and 1 mol% TR-ceramide. GUVs in panel **b** consist of 93 mol% POPC, 5 mol% Ni-NTA, 2 mol% DP-EG10-biotin and 0.1 mol% Texas Red-DHPE. Scale bar = 5 $\mu$m. **c, d** Intensity distribution along the dash line across TR-ceramide labeled GUV (**c**) and Texas Red-DHPE labeled GUV (**d**) as shown in panel **a** and **b**. Green line represents protein signal and the red line represents the lipid signal.

**Table S1**. Tubule frequency as a function of FUS LC concentration. GUVs consist of 93 mol% POPC, 5 mol% Ni-NTA, 2 mol% DP-EG10-biotin and 0.1 mol% Texas Red-DHPE. Data were collected from three independent replicates. Each replicate used a different batch of GUVs.

| Protein Conc.($\mu$M) | Experiment 1 | | | Experiment 2 | | | Experiment 3 | | | Sum | |
|---|---|---|---|---|---|---|---|---|---|---|---|
| | w/ tubule | total | Tubule frequency (%) | w/ tubule | total | Tubule frequency (%) | w/ tubule | total | Tubule frequency (%) | Avg. tubule freq. (%) | Std. dev. |
| 0.1 | 1 | 71 | 1.4 | 3 | 94 | 3.2 | 5 | 100 | 5 | 1.7 | 1.4 |
| 0.5 | 25 | 114 | 21.9 | 26 | 100 | 26 | 20 | 115 | 17.4 | 21.8 | 4.3 |
| 1 | 96 | 219 | 43.8 | 51 | 124 | 41.1 | 47 | 100 | 47 | 44 | 3.0 |
| 5 | 50 | 109 | 45.9 | 42 | 99 | 42.4 | 53 | 123 | 43.1 | 43.8 | 1.9 |
| 10 | 52 | 146 | 35.6 | 41 | 106 | 38.7 | 35 | 108 | 32.4 | 35.6 | 3.2 |
| 15 | 71 | 199 | 35.7 | 40 | 107 | 37.4 | 35 | 101 | 34.6 | 35.9 | 1.4 |
| 25 | 50 | 144 | 34.7 | 37 | 99 | 37.4 | 34 | 110 | 30.9 | 34.3 | 3.3 |

**Table S2**. Percentage of GUVs displaying protein-rich phase separation (PS) domains as a function of Ni-NTA contents and NaCl concentration. GUVs primarily consist of POPC with varying Ni-NTA, 2 mol% DP-EG10-biotin and 0.1 mol% Texas Red-DHPE. N > 100 GUVs were analyzed cumulatively from three independent replicates for each condition. Each replicate used a different batch of GUVs.

| Ni-NTA conc. (%) | NaCl Conc. ($\mu$M) | Experiment 1 | | | Experiment 2 | | | Experiment 3 | | | Sum | |
|---|---|---|---|---|---|---|---|---|---|---|---|---|
| | | PS | total | PS freq. (%) | PS | total | PS freq. (%) | PS | total | PS freq. (%) | Avg. PS freq. (%) | Std. dev. |
| | 50 | 1 | 26 | 3.8 | 2 | 27 | 7.4 | 2 | 44 | 4.5 | 5.3 | 1.9 |
| 2 | 150 | 4 | 42 | 9.5 | 6 | 43 | 14.0 | 4 | 29 | 13.8 | 12.4 | 2.5 |
| | 250 | 12 | 44 | 27.3 | 11 | 70 | 15.7 | 14 | 63 | 22.2 | 21.7 | 5.8 |
| | 50 | 7 | 48 | 14.6 | 11 | 66 | 16.7 | 6 | 35 | 17.1 | 16.1 | 1.4 |
| 5 | 150 | 26 | 100 | 26 | 34 | 124 | 27.4 | 66 | 219 | 30.1 | 27.9 | 2.1 |
| | 250 | 47 | 96 | 49.0 | 15 | 36 | 41.7 | 40 | 67 | 59.7 | 50.1 | 9.0 |
| | 50 | 23 | 36 | 63.9 | 17 | 39 | 43.6 | 34 | 63 | 54.0 | 53.8 | 10.1 |
| 15 | 150 | 23 | 73 | 31.5 | 11 | 39 | 28.2 | 43 | 112 | 38.4 | 32.7 | 5.2 |
| | 250 | 41 | 95 | 43.2 | 39 | 81 | 48.1 | 10 | 26 | 38.5 | 43.3 | 4.8 |

**Table S3**. Percentage of GUVs displaying inward tubules as a function of Ni-NTA contents and NaCl concentration. GUVs primarily consist of POPC with varying Ni-NTA, 2 mol% DP-EG10-biotin and 0.1 mol% Texas Red-DHPE. N > 100 GUVs were analyzed cumulatively from three independent replicates for each condition. Each replicate used a different batch of GUVs.

| Ni-NTA conc. (%) | NaCl Conc. ($\mu$M) | Experiment 1 | | | Experiment 2 | | | Experiment 3 | | | Sum | |
|---|---|---|---|---|---|---|---|---|---|---|---|---|
| | | Tubule | total | Tubule freq. (%) | Tubule | total | Tubule freq. (%) | Tubule | total | Tubule freq. (%) | Avg. tubule freq. (%) | Std. Dev. |
| 2 | 50 | 0 | 26 | 0 | 0 | 27 | 0 | 0 | 44 | 0 | 0 | 0 |
| | 150 | 0 | 42 | 0 | 0 | 43 | 0 | 0 | 29 | 0 | 0 | 0 |
| | 250 | 0 | 44 | 0 | 0 | 70 | 0 | 0 | 63 | 0 | 0 | 0 |
| 5 | 50 | 4 | 48 | 8.3 | 5 | 66 | 7.6 | 1 | 35 | 2.9 | 6.3 | 3.0 |
| | 150 | 47 | 100 | 47.0 | 51 | 124 | 41.1 | 96 | 219 | 43.8 | 44.0 | 2.9 |
| | 250 | 40 | 96 | 41.7 | 14 | 36 | 38.9 | 24 | 67 | 35.8 | 38.8 | 2.9 |
| 15 | 50 | 15 | 36 | 41.7 | 11 | 39 | 28.2 | 24 | 63 | 38.1 | 36.0 | 7.0 |
| | 150 | 42 | 73 | 57.5 | 20 | 39 | 51.3 | 52 | 112 | 46.4 | 51.7 | 5.6 |
| | 250 | 42 | 95 | 44.2 | 43 | 81 | 53.1 | 11 | 26 | 42.3 | 46.5 | 5.8 |

**Table S4.** Summary of GUV compositions for different membrane bending rigidity experiments

| Group | Membrane composition | Approximate Bending Rigidity from major components ($k_BT$) |
|---|---|---|
| DOPC | 83 mol% DOPC, 15 mol% Ni-NTA, 2 mol% DP-EG10 biotin, 0.1% Texas Red DHPE | 26 (10) |
| DPHPC | 83 mol% DPHPC, 15 mol% Ni-NTA, 2 mol% DP-EG10 biotin, 0.1% Texas Red DHPE | 29 (11) |
| POPC | 83 mol% POPC, 15 mol% Ni-NTA, 2 mol% DP-EG10 biotin, 0.1% Texas Red DHPE | 31 (12) |
| 1:1 POPC: Cholesterol "POPC+50% Chol" | 41.5 mol% POPC, 41.5 mol% Cholesterol, 15 mol% Ni-NTA, 2 mol% DP-EG10 biotin, 0.1% Texas Red DHPE | 37 (13) |

| | | |
|---|---|---|
| POPC+30% Sphingomyelin | 58.1 mol% POPC, 24.9 mol% Sphingomyelin, 15 mol% Ni-NTA, 2 mol% DP-EG10 biotin, 0.1% Texas Red DHPE | 42 (14) |
| 1:1 Sphingomyelin: Cholesterol "SM+50% Chol" | 41.5 mol% Sphingomyelin, 41.5 mol% Cholesterol, 15 mol% Ni-NTA, 2 mol% DP-EG10 biotin, 0.1% Texas Red DHPE | 76 (15) |

In addition, a batch of GUVs labeled with TR-Ceramide was made as a comparison. The composition was 83 mol% POPC, 15 mol% Ni-NTA, 2 mol% DP-EG10-biotin and 1 mol% TR-Ceramide.

**Supplementary Movie 1.** Confocal image series of a GUV containing protein-lined tubules formed upon exposure to his-FUS LC (1 μM). Tubules are flexible and display dynamic changes in morphology. The images were taken along an axis perpendicular to the imaging plane. The images are 0.5 μm apart. The composition of the GUV was 83 mol% POPC, 15 mol% DOGS Ni-NTA, 2 mol% DP-EG10-biotin, and 0.1 mol% Texas Red-DHPE. The buffer composition was 25 mM HEPES, 150 mM NaCl, pH 7.4. The scale bar is 5 μm long.

**Supplementary Movie 2.** On the top surface of a giant unilamellar vesicle (GUV), domains of the protein-depleted phase (dark), move randomly within the protein-enriched phase (bright), confocal image series. The ability of the depleted phase regions to move rapidly within the enriched phase, as well as fluctuations in the boundaries of the depleted phase, suggest that the enriched phase is liquid-like, rather than a rigid solid. The his-FUS LC protein was labeled with Atto-488. The protein concentration was 5 μM. The composition of the GUVs was 83 mol% POPC, 15 mol% DOPG-Ni-NTA, 2 mol% DP-EG10-Biotin, 0.1% Texas Red DHPE and the experiment was performed in 25 mM HEPES, 150 mM NaCL, pH 7.4 buffer. The scale bar is 5 μm long.

### Materials and Methods

#### Reagents

1,2-dioleoyl-sn-glycero-3-phosphocholine (DOPC), 1,2-diphytanoyl-sn-glycero-3-phosphocholine (DPHPC), 1-palmitoyl-2-oleoyl-glycero-3-phosphocholine (POPC), Sphingomyelin (Brain, Porcine), cholesterol and 1,2-dioleoyl-sn-glycero-3-[(N-(5-amino-1-carboxypentyl)iminodiacetic acid)succinyl] (nickel salt) (DOGS-NTA-Ni), were purchased from Avanti Polar Lipids, Inc. NaCl, Tris hydrochloride (TrisHCl), 4-(2-hydroxyethyl)-1-piperazineethanesulfonic acid (HEPES), neutravidin, Texas Red-DHPE, isopropyl-β-D-thiogalactopyranoside (IPTG), β-mercaptoethanol (β-ME), BODIPY™ TR Ceramide and Triton X-100 were obtained from Thermo Fisher Scientific. 2-(N-Morpholino)ethanesulfonic acid hydrate, 4-Morpholineethanesulfonic acid (MES hydrate), 3-(Cyclohexylamino)-1-propanesulfonic acid (CAPS), Urea, $NaH_2PO_4$, $Na_2HPO_4$, $Na_3PO_4$, sodium tetraborate, Ethylenediaminetetraacetic acid (EDTA), phenylmethanesulfonyl fluoride (PMSF), EDTA-free protease inhibitor tablets, imidazole, poly-L-lysine (PLL), and ATTO-488 NHS-ester were purchased from Sigma-Aldrich. Dipalmitoyl-decaethylene glycol-biotin (DP-EG10-biotin) was kindly provided by Darryl Sasaki from Sandia National Laboratories, Livermore, CA (1). Amine reactive PEG (mPEG-Succinimidyl Valerate MW 5000) and PEG-biotin (Biotin-PEG SVA, MW 5000) were purchased from Laysan Bio, Inc. Fetal bovine serum (FBS), trypsin, penicillin, streptomycin, L-glutamine, phosphate-buffered saline (PBS), Ham's F-12, Ham's F-12 without phenol red, Dulbecco's modified Eagle medium (DMEM) and DMEM without phenol red were purchased from GE Healthcare. N-ethylmalemide (NEM) was purchased from Bio Basic. All reagents were used without further purification.

**Plasmids**

The pRP1B FUS 1-163 plasmid for FUS LC (residues 1-163) protein incorporating a TEV cleavable N-terminal hexahistidine tag was a gift from Nicolas Fawzi Lab, Brown University (2). This plasmid is available from AddGene (https://www.addgene.org/127192/). The plasmid for expression of GFP-FUS LC (residues 2-214) was generously provided by the lab of Steven McKnight at the University of Texas Southwestern Medical Center (3). The pRSET vector coding for the nondimerizable hexa-his-tagged eGFP (hisGFP A206K) was kindly shared by Dr. Adam Arkin (University of California, Berkeley). The plasmids for hnRNPA2 LC and LAF-1 RGG domain were also obtained from AddGene (https://www.addgene.org/98657/ and http://www.addgene.org/124929/, respectively).The plasmid for mammalian expression of TfR-Δecto-BFP-HA-GFPnb was generated by inserting a hemagglutinin (HA)-tag into the TfR-Δecto-BFP-GFPnb plasmid previously described (4). First, the GFPnb sequence from this original plasmid was amplified by polymerase chain reaction (PCR) using primers that introduced the HA-tag sequence. The amplified HA-GFPnb sequence was then restriction cloned back into the original plasmid using NotI sites. All constructs were confirmed by DNA sequencing. Notably, the HA tag was included for screening purposes and did not play a functional role in the present study.

**Production of stable cell line and cell culture**

Human retinal pigmented epithelial (RPE) cells (ARPE-19) were purchased from American Type Culture Collection (Manassas, VA). RPE cells were stably transfected with the plasmid encoding TfR-Δecto-BFP-HA-GFPnb via lentiviral transfection. The TfR-Δecto-BFP-HA-GFPnb plasmid described above was subcloned onto the pLJM1 viral transfer vector (Addgene 19319).To generate lentiviruses, human embryonic kidney (HEK) 293T cells (ATCC) were co-transfected with the transfer plasmid, packaging plasmid Δ8.9, and the envelope plasmid VSVG using FuGENE transfection reagent (Promega). The transfected HEK 293T cells were incubated at 37ºC for 48 hours, after which the virus-containing media was collected and filtered through a 0.45 µm average pore sized filter. For transduction, the filtered virus was then added to RPE cells with 8 µg/mL Polybrene. The transduced RPE cells were selected with 2 µg/mL puromycin for 7 days, before cell sorting was performed by flow cytometry to select for cells containing BFP fluorescence. After selection, these cells were cultured in 1:1 F12:DMEM supplemented with 10% FBS, 20 mM HEPES, and 1% penicillin, 1% streptomycin, 1% L-glutamine (PSLG). Cells were incubated at 37 °C with 5% $CO_2$ and passaged every 48−72 hours.

**Protein expression and purification**

Expression and purification of his-FUS LC was carried out according to previous report (5), with several modifications. In brief, his-FUS LC was overexpressed in *E. Coli* BL21(DE3) cells. Pellets of cells expressing his-FUS LC were harvested from 1 liter cultures induced with 1 mM IPTG after 4-hour incubation at 37°C and 220 RPM when OD 600 was around 0.8. The pellets were then lysed in a buffer containing 0.5 M TrisHCl pH 8, 5 mM EDTA, 5% glycerol, 10mM β-ME, 1mM PMSF, 1% Triton X-100 and one EDTA-free protease inhibitor tablet (Sigma Aldrich) for 5 min on ice and then sonicated. The cell lysates were centrifuged at 40,000 RPM for 40 min and his-FUS LC resided in the insoluble fraction after centrifugation. Therefore, the insoluble fraction was resuspended in 8M urea, 20 mM NaPi pH 7.4, 300 mM NaCl and 10 mM imidazole. The resuspended sample was then centrifuged at 40,000 RPM for 40 min. In denaturing conditions, his-FUS LC is urea-soluble and so at this point resided in the supernatant. This supernatant was then mixed with Ni-NTA resin (G Biosciences, USA) for 1 hour at 4°C. The Ni-NTA resin was settled in a glass column and washed with the above solubilizing buffer. The bound proteins were eluted from the Ni-NTA resin with a buffer containing 8M urea, 20 mM NaPi pH 7.4, 300 mM NaCl and 500 mM imidazole. Unlike previous described applications, the TEV-cleavable his-tag of this protein was not removed, as it was needed for membrane binding. The purified proteins were then buffer-exchanged into 20 mM CAPS pH 11 storage buffer using 3K Amicon Ultra centrifugal filters (Millipore, USA). Small aliquots of the protein were frozen in liquid nitrogen at a protein concentration of approximately 1 mM.

His-hnRNPA2 LC purification was performed as described previously (6). When expressed in bacteria, his-hnRNPA2 LC is found in inclusion bodies. The expression and purification protocol for his-hnRNPA2 is the same as the protocol for purifying his-FUS

LC, except that his-hnRNPA2 LC was finally exchanged into the storage buffer containing 20 mM MES, 8 M Urea pH 5.5.

Expression and purification of his-LAF-1 RGG was performed as described previously (7). Briefly, his-LAF-1 RGG was overexpressed in *E. Coli* BL21(DE3) cells. Pellets of cells expressing his-FUS LC were harvested from 1 liter cultures induced with 0.5 mM IPTG after overnight incubation at 18°C and 220 RPM when OD 600 was around 0.8. The pellets were then lysed in a buffer containing 20 mM Tris, 500mM NaCl, 20 mM imidazole, and one EDTA-free protease inhibitor tablet (Sigma Aldrich) for 5 min on ice and then sonicated. The cell lysates were centrifuged at 40,000 RPM for 40 min and his-LAF-1 RGG was in the lysate after centrifugation. The lysate was then mixed with Ni-NTA resin (G Biosciences, USA) for 1 hour. The Ni-NTA resin was settled in a glass column and washed with a buffer containing 20mM Tris, 500 mM NaCl, 20 mM imidazole, at pH 7.5. The bound proteins were eluted from the Ni-NTA resin with a buffer containing 20 mM Tris, 500 mM NaCl, 500 mM imidazole, at pH 7.5. The purified proteins were then buffer-exchanged into a buffer containing 20mM Tris, 500mM NaCl, at pH 7.5. Small aliquots of the protein were frozen in liquid nitrogen at a protein concentration of approximately 120 μM and stored at -80 °C. To promote solubility of the protein, the entire purification process was performed at room temperature, except for the cell lysis process, which was done on ice.

Expression and purification of GFP-FUS LC was also conducted based on previously reported protocols (3). Briefly, pellets of *E. Coli* BL21(DE3) cells expressing GFP-FUS LC were harvested from 1 liter cultures induced with 1 mM IPTG after overnight incubation at 16°C and 220 RPM. The cell pellets were then lysed in a buffer containing 50 mM Tris-HCl pH7.5, 500 mM NaCl, 20 mM BME, 1mM PMSF, 1% Triton X-100 and one protease inhibitor tablet for 5 min on ice, and then sonicated. The cell lysates were centrifuged at 40,000 RPM for 40 min to separate the lysate from the insoluble fraction and GFP-FUS LC resided in the supernatant after centrifugation. Therefore, the supernatant was then mixed with Ni-NTA resin for 2 hours at 4°C. The Ni-NTA resin was then packed in a glass column and washed with a buffer containing 20 mM $Na_3PO_4$ and 150mM NaCl pH 7.4, 20 mM imidazole, 20 mM BME, and 0.1 mM PMSF. The bound protein was eluted from the Ni-NTA resin with a buffer containing 20 mM $Na_3PO_4$ and 150mM NaCl pH 7.4, 500 mM imidazole, 20 mM BME, and 0.1 mM PMSF. The buffer was then exchanged into a storage buffer consisting of 20mM $Na_3PO_4$ pH 7.4 buffer using 10K Amicon Ultra centrifugal filters. The purified proteins were further concentrated and flash frozen in small aliquots at a concentration of approximately 120mM.

His-GFP was used as a controlled protein and its purification was performed according to a previously published protocol (8).

**Protein labeling**
His-FUS LC was labeled with Atto-488, an amine-reactive, NHS ester-functionalized green dye. Labeling occurred at or near the N-terminus because only the N-terminus and a lysine at residue position 5 in the leader sequence preceding the N-terminal region hexa-histidine tag were expected to react with the NHS-functionalized dye. The labeling reaction took place in a 50mM HEPES buffer at pH 7.4. Dye was added to the protein in 2-fold stoichiometric excess and allowed to react for 30 min at room temperature, empirically resulting in labelling ratio near 1:1 dye: protein. Labeled protein was then buffer-exchanged into 20mM CAPS pH 11 buffer and separated from unconjugated dye using 3K Amicon columns. His-hnRNPA2 LC and his-LAF-1 RGG were also labeled with Atto-488 following the same process in corresponding buffers. His-hnRNPA2 LC was buffer-exchanged into 20 mM MES, pH 5.5 buffer prior to reacting with the dye, and then buffer-exchanged back into 20 mM MES, 8 M Urea pH 5.5 for storage. His-LAF-1 RGG was labeled in its storage buffer (20 mM Tris, 500 mM NaCl, pH 7.5). Protein and dye concentrations were monitored using UV–Vis spectroscopy. Labeled proteins were dispensed into small aliquots, flash frozen in liquid nitrogen and stored at -80°C. For all experiments involving labeled protein, a mix of 90% unlabeled / 10% labeled protein was used.

**GUV preparation**
For all GUV experiments except for experiments with different GUV membrane bending rigidity, GUVs were made of POPC, Ni-NTA, 2 mol% DP-EG10 biotin and an additional 0.1 mol% Texas Red DHPE lipids, with the contents of POPC and Ni-NTA adjusted accordingly.

For experiments with different GUV membrane bending rigidity, GUVs of six different compositions were prepared based on previous reports. The detailed compositions are shown in Table S4, above.

GUVs were prepared according to published protocols (9). Briefly, lipid mixtures dissolved in chloroform were spread into a film on indium-tin-oxide (ITO) coated glass slides (resistance ~8-12W sq$^{-1}$) and further dried in a vacuum desiccator for at least 2 h to remove all of the solvent. Electroformation was performed at 55°C in glucose solution. 360 milliosmole glucose solution was employed for making GUVs, except for GUVs used in different salt concentration experiments, where 560 milliosmole glucose solution was adopted to have more capacity to modulate osmolarity. The voltage was increased every 3 min from 50 to 1400 mVpp for the first 30 min at a frequency of 10 Hz. The voltage was then held at 1400 mVpp for 120 min and finally was increased to 2200 mVpp for the last

30 min during which the frequency was adjusted to 5 Hz. GUVs were stored in 4°C and used within 1d after electroformation.

**GUV tethering**

Prior to tethering, the osmolality of the GUV solution and experimental buffers were measured using a vapor pressure osmometer (Wescor). GUVs were tethered to glass coverslips as previously described (16). Briefly, glass cover slips were passivated with a layer of biotinylated PLL-PEG, using 5 kDa PEG chains. GUVs doped with 2 mol% DP-EG10-biotin were then tethered to the passivated surface using neutravidin.

PLL-PEG was synthesized by combining amine reactive PEG and PEG-biotin in molar ratios of 98% and 2%, respectively. This PEG mixture was added to a 20 mg/mL mixture of PLL in a buffer consisting of 50mM sodium tetraborate (pH 8.5), such that the molar ratio of lysine subunits to PEG was 5:1. The mixture was continuously stirred at room temperature for 6 h and then buffer exchanged into 25mM HEPES, 150mM NaCl (pH 7.4) using Zeba$^{TM}$ Spin Desalting Column (ThermoFisher Scientific).

Buffer used for dilution and rinsing was 25mM HEPES, 150 mM NaCl pH 7.4 buffer unless specifically stated. For GUV experiments under different salt concentrations, 25mM HEPES pH 7.4 containing 50mM, 150mM, 250mM NaCl were used, respectively, to achieve salt gradient. Osmolarity balance was maintained by the addition of glucose to the buffer.

Imaging wells consisted of 5 mm diameter holes in 0.8 mm thick silicone gaskets (Grace Bio-Labs). Gaskets were placed directly onto no.1.5 glass coverslips (VWR International), creating a temporary water-proof seal. Prior to well assembly, gaskets and cover slips were cleaned in 2% v/v Hellmanex III (Hellma Analytics) solution, rinsed thoroughly with water, and dried under a nitrogen stream. In each dry imaging well, 20 µL of PLL-PEG was added. After 20 min of incubation, wells were serially rinsed with appropriate buffer by gently pipetting until a 15,000-fold dilution was achieved. Next, 4 µg of neutravidin dissolved in 25mM HEPES, 150mM NaCl (pH 7.4) was added to each sample well and allowed to incubate for 10 min. Wells were then rinsed with the appropriate buffer to remove excess neutravidin. GUVs were diluted in appropriate buffer at ratio of 1:13 and then 20 µL of diluted GUVs was added to the well and allowed to incubate for 10 min. Excess GUVs were then rinsed from the well using the appropriate buffer and the sample was subsequently imaged using confocal fluorescence microscopy.

**GUV fluorescence imaging**

Imaging experiments were performed using a spinning disc confocal super resolution microscope (SpinSR10, Olympus, USA) equipped with a 1.49 NA/100X oil immersion objective. Laser wavelengths of 488 and 561 nm were used for excitation. Image stacks taken at fixed distances perpendicular to the membrane plane (0.5 μm steps) were acquired immediately after GUV tethering and again after protein addition. At least 30 fields of views were randomly selected for each sample for further analysis prior to and after the addition of protein, respectively.

**Giant Plasma Membrane Vesicles (GPMVs)**

GPMVs were derived from RPE cells that stably expressed TfR-Δecto-BFP-HA-GFPnb receptor, according to published protocols (17).These donor cells were grown in 100 × 20 mm culture dishes. Prior to extraction of GPMVs, these cells were rinsed twice with 2 mL GPMV buffer (10 mM HEPES, 2 mM $CaCl_2$, 150 mM NaCl, pH 7.4) and once with 2 mL active buffer (GPMV buffer containing 2mM N-ethylmalemide). The cells were then incubated in 4 mL active buffer at 37 °C for 10 h. During this time, GPMVs budded spontaneously from the plasma membrane of the cells. Following this incubation, the buffer containing GPMVs was collected and spun at 300 x g for 3 min at 4 °C to remove any detached cells. The supernatant was transferred to a fresh tube and centrifuged at 17,000 x g for 23 min at 4°C. The GPMV pellet was then resuspended in 200 μL fresh GPMV buffer and used immediately for imaging experiments.

**GPMV imaging**

Similar to imaging experiments with GUVs, 20 μL of GPMV-containing solution was added onto a glass coverslip. GPMVs were allowed to settle onto the coverslip surface at 4°C for 10 min. A spinning disc confocal microscope (Zeiss Axio Observer Z1 with Yokagawa CSU-X1M) was used to image GPMVs. GFP-FUS LC protein diluted in GPMV buffer was added to the sample at a 2 μM final concentration. GFP alone at the same concentration was adopted as a control protein that does not form condensates. Image stacks taken perpendicularly to the coverslips (0.5 μm steps) were acquired and at least 30 fields of views were randomly selected for each sample for further analysis prior to and after the addition of protein, respectively.

**Statistical analysis**

All GPMV and GUV experiments were repeated 3 times for each condition reported. ImageJ was employed to analyze confocal images. At least 100 GPMVs and GUVs were examined under each condition. The diameter of each tubule was determined by drawing a line perpendicular to the tubule at three different places along its length and calculating the average diameter. The distribution of tubule diameters for each condition was derived from measurements of at least 100 vesicles. To assess the significance of comparisons between conditions, an unpaired t-test was performed. Error bars in graphs represent either standard error or standard deviation as stated in figure captions.

### Model development

#### Assumptions

- We treat the lipid bilayer as an elastic shell assuming that the thickness of the bilayer is negligible compared to the radii of the membrane curvatures (18). This assumption allows us to model the bending energy of the membrane using the Helfrich–Canham energy (19, 20).
- We assume that the membrane is locally incompressible (21) using a Lagrange multiplier to implement this constraint (22, 23).
- We ignore inertia and assume that the membrane is at mechanical equilibrium at all times (24-26).
- We model the net effect of the phase separation of the proteins on the membrane surface by spontaneous curvature ( $C$ ). The magnitude of the spontaneous curvature represents the curvature-generating capability of the protein domain to bend the membrane (27, 28) and the induced area difference between the leaflets (29-31).
- We assume that the protein phase separation induces a spherical membrane cap shape with a constant curvature (Fig. 3c) (32) for ease of computation.
- To keep the mathematics tractable, we assume that the geometry is rotationally symmetric (see Fig. 3c) (20, 25, 33). This allows us to obtain solutions for different morphologies of the membrane tubules with a relatively simple numerical calculation.

#### Helfrich energy and mechanical equilibrium

For the local energy density of a lipid bilayer membrane, we use the modified version of the Helfrich energy including the spatially varying spontaneous curvature and bending modulus as (28, 33-35).

$$W = \kappa(\theta^{\xi})\left(H - C(\theta^{\xi})\right)^2 + \kappa_G K \qquad \text{(S1)}$$

where $W$ is the local energy density, $H$ is the mean curvature, $K$ is the Guassian curvature, $C$ is the induced spontaneous curvature due to any asymmetry between the leaflets, $\kappa$ is the bending modulus, $\kappa_G$ is the Gaussian modulus, and $\theta^{\xi}$ is a representation of surface coordinates where $\xi \in [1, 2]$ . It should be noted that Eq. (S1) is different from the standard Helfrich energy by a factor of 2. We take this net effect into consideration by choosing the value of the bending modulus to be twice that of the standard value of bending modulus typically used for lipid bilayers (19). The induced spontaneous curvature ($C$) in Eq. S1 can be related to the area difference between the bilayer leaflets given in the literature (29-31).

$$\Delta A - \Delta A_0 \equiv \frac{4d\kappa CA}{\pi\kappa_r} \tag{S2}$$

where $d$ is the lipid bilayer with thickness, $\kappa_r$ is the nonlocal membrane bending modulus, and $A$ is the total surface area of the neutral plane. $\Delta A$ and $\Delta A_0$ are the relaxed initial and bent area differences between the membrane leaflets, respectively. Thus, induced spontaneous and area difference are not independent parameters. Any stationary shape of membrane that is a result of area difference elasticity model (ADE) therefore can be also modeled by the spontaneous curvature given in Eq. S2.

As we showed previously, the normal variation of total energy of the membrane gives the so-called "shape equation," (25, 28, 34)

$$\underbrace{\Delta[\kappa(H-C)] + 2\kappa(H-C)(2H^2-K) - 2\kappa H(H-C)^2}_{\text{Elastic effects}} = \underbrace{p + 2\lambda H}_{\text{Capillary effects}} \tag{S3}$$

Where $\Delta$ is the surface Laplacian, $p$ is the pressure difference across the membrane, and $\lambda$ can be interpreted to be the membrane tension (28).

A consequence of spatial variation of membrane properties and protein density is that $\lambda$ is not homogeneous along the membrane (25, 28, 36). Therefore, the balance of forces tangential to the membrane gives the spatial variation of membrane tension as,

$$\underbrace{\nabla\lambda}_{\substack{\text{Gradient of}\\ \text{surface tension}}} = \underbrace{2[\kappa(H-C)]\frac{\partial C}{\partial\theta^{\xi}}}_{\text{Protein density variation}} - \underbrace{\frac{\partial\kappa}{\partial\theta^{\xi}}(H-C)^2}_{\substack{\text{Bending modulus–induced}\\ \text{variation}}} \tag{S4}$$

where $\nabla$ is the partial derivative with respect to the coordinate ($\theta^{\xi}$).

**Parametrization in axisymmetric coordinates**

We define a surface of revolution (Fig. 3c) by

$$\boldsymbol{r}(s,\theta) = R(s)\boldsymbol{e_r}(\theta) + Z(s)\mathbf{k} \tag{S5}$$

where ($\boldsymbol{e_r}$, $\boldsymbol{e_\theta}$, $\mathbf{k}$) is the basis coordinate, $s$ is the arc length along the curve, $R(s)$ is the radius from the axis of rotation and $Z(s)$ is the height from the base plane. We $\psi$ as the angle made by the tangent with respect to the vertical such that $R' = \cos\psi$ and $Z' = \sin\psi$, which satisfies the identity $(R')^2 + (Z')^2 = 1$. Here, $()'$ is the partial derivative with respect to the arc length. We can define the normal and tangent vectors to the surface as

$$\boldsymbol{n} = -\sin\psi\,\boldsymbol{e_r}(\theta) + \cos\psi\,\boldsymbol{k},\ \ \boldsymbol{a_s} = \cos\psi\,\boldsymbol{e_r}(\theta) + \sin\psi\ \boldsymbol{k} \tag{S6}$$

The tangential ($\kappa_\nu$) and transverse ($\kappa_\tau$ ) curvatures are given by

$$\kappa_\nu = \psi', \qquad \kappa_\tau = sin\,\psi/R \tag{S7}$$

and the mean curvature ($H$) and the Gaussian curvature ($K$) are defined as

$$H = \frac{1}{2}(\kappa_\nu + \kappa_\tau) = \frac{1}{2}(\psi' + sin\,\psi/R) \tag{S8}$$

To simplify the governing equations to first- order differential equations, we define $L = \frac{1}{2\kappa}R(W_H)'$ giving us (25, 34, 35),

$$R' = \cos\psi,\ Z' = \sin\psi,\ R\psi' = 2RH - \sin\psi,\ RH' = L + RC' - \frac{R\kappa'}{\kappa}(H - C),$$
$$\frac{L\prime}{R} = \frac{p}{\kappa} + 2H\left[(H-C)^2 + \frac{\lambda}{\kappa}\right] - 2(H-C)\left[H^2 + \left(H - \frac{\sin\psi}{R}\right)^2\right] - \frac{\kappa' L}{\kappa R}, \tag{S9}$$
$$\lambda' = 2[\kappa(H-C)]C' - \kappa'(H-C)^2.$$

In asymmetric coordinates, the total area of the manifold ($A$) can be expressed in term of arc length as,

$$A(s) = 2\pi\int_0^S R(\eta)d\eta \rightarrow \frac{dA}{ds} = 2\pi R. \tag{S10}$$

Thus, Eq. (S10) allows us to convert the system of equations in Eq. (S9) that is in term of arc length to a system in terms of area given by

$$2\pi R\dot{R} = \cos\psi, 2\pi R\,\dot{Z} = \sin\psi, 2\pi R^2\dot{\psi} = 2RH - \sin\psi, 2\pi R^2\dot{H} = L + 2\pi R^2(\dot{C} - \frac{\dot{\kappa}}{\kappa}(H - C)),$$
$$2\pi\dot{L} = \frac{p}{\kappa} + 2H\left[(H-C)^2 + \frac{\lambda}{\kappa}\right] - 2(H-C)\left[H^2 + \left(H - \frac{\sin\psi}{R}\right)^2\right] - 2\pi\frac{\dot{\kappa}L}{\kappa}, \tag{S11}$$
$$2\pi R\dot{\lambda} = 4\pi R[\kappa(H-C)]\dot{C} - 2\pi R\dot{\kappa}(H-C)^2.$$

where ( $\dot{}$ ) denotes the derivative with respect to the area.

To non-dimensionalize the system of equations (Eq. (S11)), we use two parameters, the radius of the GUV ($R_0$), and lipid bilayer bending modulus ($\kappa_0$). Using these constants, we can define

$$\zeta = \frac{A}{2\pi R_0^2},\ y = \frac{Z}{R_0},\ x = \frac{R}{R_0},\ h = HR_0,\ c = CR_0,\ l = LR_0,$$
$$\tilde{\lambda} = \frac{\lambda R_0^2}{\kappa_0},\ G = KR_0^2,\ \tilde{p} = \frac{pR_0^2}{\kappa_0},\ \widetilde{\kappa_G} = \frac{\kappa_G}{\kappa_0},\ \tilde{\kappa} = \frac{\kappa}{\kappa_0}. \tag{S12}$$

Rewriting Eq. (S11) in terms of the dimensionless variables in Eq. (S12), we get (25)

$$x\dot{x} = \cos\psi,\ x\dot{y} = \sin\psi,\ x^2\dot{\psi} = 2xh - \sin\psi,\ x^2\dot{h} = l + x^2(\dot{c} - \frac{\dot{\tilde{\kappa}}}{\tilde{\kappa}}(h - c)),$$
$$\dot{l} = \frac{\tilde{p}}{\tilde{\kappa}} + 2h\left[(h-c)^2 + \frac{\tilde{\lambda}}{\tilde{\kappa}}\right] - 2(h-c)\left[h^2 + \left(h - \frac{\sin\psi}{x}\right)^2\right] - \frac{\dot{\tilde{\kappa}}l}{\tilde{\kappa}} \tag{S13}$$

$\dot{\tilde{\lambda}} = 2[\tilde{\kappa}(h-c)]\dot{c} - \dot{\kappa}(h-c)^2.$

**Boundary conditions**

In order to solve the system of equations (Eq. S9), we need to provide six boundary conditions. We consider an axisymmetric circular patch of membrane. To impose the continuity at $s = 0$, we require to set $\psi = \pi/2$ and $R = R_b$ where we used the suggested length scale in Naito et al. (37) to relate the radius of undoloid to the physical properties of the membrane. At the other boundary far from the center of the patch ($s = s_{max}$), we set $Z = 0$ to be sure the membrane does not lift off. We also assume that the membrane at the far boundary remains flat, so we set $\psi = \psi' = 0$ and prescribe the tension as $\lambda = \lambda_0$. The boundary conditions can be summarized as

$$R(0) = R_b,\ \psi(0) = \frac{\pi}{2},\ Z(s_{max}) = 0, \psi(s_{max}) = 0,\ \psi'(s_{max}) = 0,\ \lambda(s_{max}) = \lambda_0. \quad \text{(S14)}$$

**Delaunay shapes and other solution of Helfrich energy minimization in axisymmetric coordinates**

The general shape equation (Eq. S3) is a nonlinear fourth order differential equation. Delaunay's surface including catenoids, unduloids, circular cylinder, and spheres are surfaces of revolution with constant mean curvature and they are found to be a solution of the Helfrich shape equation (38, 39). The general equation describing the Delaunay's shape is given by

$$sin\,\psi(R) = aR + \frac{d}{R}, \quad \text{(S15)}$$

where $a$ and $d$ are constants determining the type of surfaces. For example, if *(i)* $a = 0$, Eq. S15 gives the catenoid shapes, if *(ii)* $0 < ad < 1/4$, Eq. S15 gives the unduloid shapes, and if *(iii)* $ad < 0$, Eq. S15 corresponds to the nodoid surfaces (37, 40). It was shown (37, 41) that the Delaunay shapes can be extended as

$$sin\,\psi(R) = aR + b + \frac{d}{R}, \quad \text{(S16)}$$

where $b$ is a constant and for the special case of $b = 0$, Eq. S16 reduces to Delaunay surfaces in Eq.S15. By substituting Eq. S16 in the Helfrich shape equation (Eq. S3) we can find the constant $a$, $b$, and $d$ as the function of the physical properties of the membrane as

$$2aC_0 = \frac{\lambda}{\kappa} + C_0^2, b = \pm\sqrt{2 + 4(\frac{a}{2C_0} - 1)},\ d = \frac{1}{2C_0}. \quad \text{(S17)}$$

It should be mentioned that in our Helfrich equation (Eq.S1), bending rigidity ($\kappa$) is two times larger than the given bending rigidity in Naito et al. (37). Also, in our Helfrich equation (Eq.S1), the spontaneous curvature is a representation of any induced asymmetry in the mean curvature ($H$) while in Naito et al., the spontaneous curvature refers to the total curvature. To convert the given equations in Naito et al. (37) to our parametrization, we multiplied the given spontaneous curvature in (37) by two and divide the bending rigidity by two.

Substituting Eq. S17 into Eq. S15 and using the length scale $R_m = \frac{1}{\sqrt{2aC_0}}$ we get (37)

$$sin\,\psi(r) = \alpha(r + r^{-1}) - \sqrt{4\alpha^2 - 2}, \tag{S18}$$

where $\alpha = (2R_m C_0)^{-1}$ and $r = \frac{R}{R_m}$ . It should be mentioned that in Eq.S18, we just considered the negative branch of b due to the constraint of $|sin\,\psi| \leq 1$. Also, to satisfy the same constrain, $\alpha \leq 0.75$ . In Fig. 3a, we plotted the unduloid like shapes corresponding to Eq. S17 for two values of $\alpha$. For $\alpha \to 0.75$, the unduloidlike shape becomes like a circular cylinder and with increasing $\alpha$, multiple spheres form along the unduloid similar to a string of pearls. In addition to Delaunay shapes, there are multiple analytical studies that have shown how the curved proteins can bend the membrane into the pearled-shaped structures (42-45).

**Numerical calculation**

We solved the system of first-order differential equations using 'bvp4c' solver in MATLAB. Here, we summarize the steps and assumptions that we used the simulation.

- All the simulations were performed for a fixed area of the membrane.
- The mesh size on the domain was chosen such that it was (initially) small around the s = 0 and then increased by moving toward the far away boundary ($s = s_{max}$)
- To ensure a smooth transition in the induced spontaneous curvature ($C$) along the tubular domain, we assumed that the spontaneous curvature decreases linearly (46) using a hyperbolic tangent function to define it as

$$C = \frac{C_0}{2}\left(\frac{A - A_0}{A_0}\right)[\tanh(g(A - A_0))], \tag{S19}$$

where $A_0$ represents the area of the phase-separated protein and $g$ is a constant. You can find a more detailed analysis of the choice of the spontaneous curvature function and the parametric sensitivity of the shape equation in (47).

**Table S4**. Value of parameters in simulation

| Figures | Parameters and values |
|---|---|
| Figure 3d | $C_0 = 1.34 - 3.5 \quad \mu m^{-1}$ |

| | |
|---|---|
| | $\kappa = 22 - 85\ k_B T$<br>$\lambda_0 = 0.9\ pN/\mu m$<br>$\kappa_{\text{ratio}} = 1$<br>$A_0 = 15\ \mu m^2$ |
| Figure 3f | $C_0 = 3.5\ \mu m^{-1}$<br>$\kappa = 22 - 85\ k_B T$<br>$\lambda_0 = 0.9\ pN/\mu m$<br>$\kappa_{\text{ratio}} = 20$<br>$A_0 = 15\ \mu m^2$ |

**Table S5**. Notation used in the model

| Notation | Description | Units |
|---|---|---|
| $W$ | Local energy per unit area | $pN/\mu m$ |
| H | Mean curvature of surface | $\mu m^{-1}$ |
| K | Gaussian curvature of surface | $\mu m^{-1}$ |
| C | Spontaneous curvature | $\mu m^{-1}$ |
| $\kappa$ | Bending modulus | $k_B T$ |
| $\kappa_G$ | Gaussian modulus | $k_B T$ |
| $\theta^\xi$ | Surface coordinate | |
| $\boldsymbol{\kappa_\tau}$ | Transverse curvature | $\mu m^{-1}$ |
| $\boldsymbol{\kappa_\nu}$ | Tangential curvature | $\mu m^{-1}$ |
| $\lambda$ | Membrane tension | $pN/\mu m$ |
| $\psi$ | Angle between radial and tangential vectors | |
| $\boldsymbol{n}$ | Normal vector to membrane surface | Unit vector |
| $\boldsymbol{a_s}$ | Tangential vector to membrane surface | Unit vector |
| $R$ | Radial distance | $\mu m$ |
| $Z$ | Elevation from base plane | $\mu m$ |
| $s$ | Arc length | $\mu m$ |
| $A$ | Membrane area | $\mu m^2$ |
| $A_0$ | Area of the protein domain | $\mu m^2$ |
| $p$ | Pressure difference across the membrane | $pN/\mu m^2$ |
| $L$ | Shape equation variable | $\mu m^{-1}$ |
| $\boldsymbol{k}$ | Altitudinal basis vector | Unit vector |
| $\boldsymbol{e}(\theta)$ | Azimuthal basis vector | Unit vector |
| $\boldsymbol{e_r}(\theta)$ | Radial basis vector | Unit vector |
| $x$ | Dimensionless radial distance | |
| $y$ | Dimensionless height | |
| $h$ | Dimensionless mean curvature | |

| | |
|---|---|
| $c$ | Dimensionless spontaneous curvature |
| $l$ | Dimensionless $L$ |
| $\tilde{\lambda}$ | Dimensionless membrane tension |
| $\tilde{\kappa}$ | Dimensionless bending modulus |
| $\tilde{\kappa}_G$ | Dimensionless Gaussian modulus |

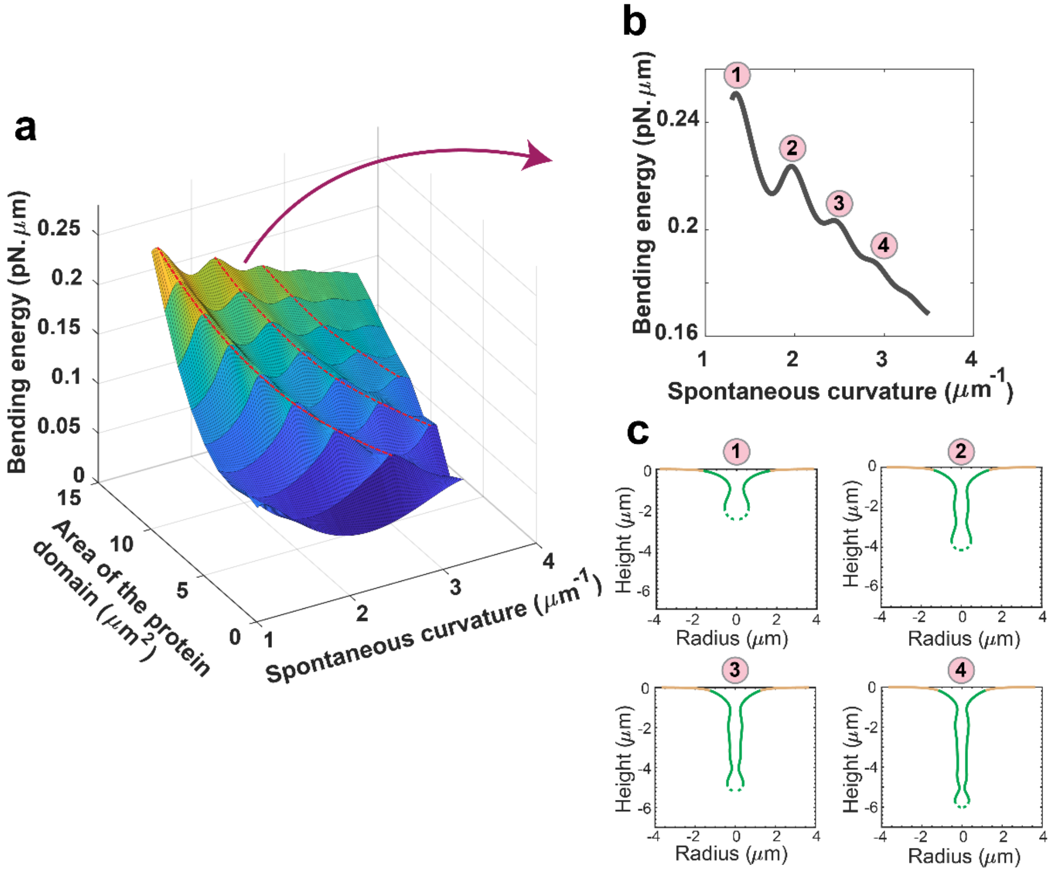


**Figure S3**. Spontaneous curvature and the area of the protein domain regulate the morphology of the undulated tubules. (**a**) Bending energy of the protein domain as the function of the spontaneous curvature and area of the protein domain. For a small area of the protein domain, there is just a local maximum with increasing the magnitude of the spontaneous curvature. However, for the larger area of the protein domain, the bending energy has an oscillation behavior with multiple local maxima as a function the spontaneous curvature. With decreasing the area of the protein domain, the bending energy decreases and also local maximum shifts toward the larger values of the spontaneous curvature (red dotted lines). (**b**) The bending energy as the function of spontaneous curvature for $A_0 = 15\ \mu m^2$. With increasing the magnitude of spontaneous curvature, there are multiple local maxima. Here, we labeled the four local maximum with numbers one to four. (**c**) The morphology of the membrane at the local maximum bending energy points in panel b.

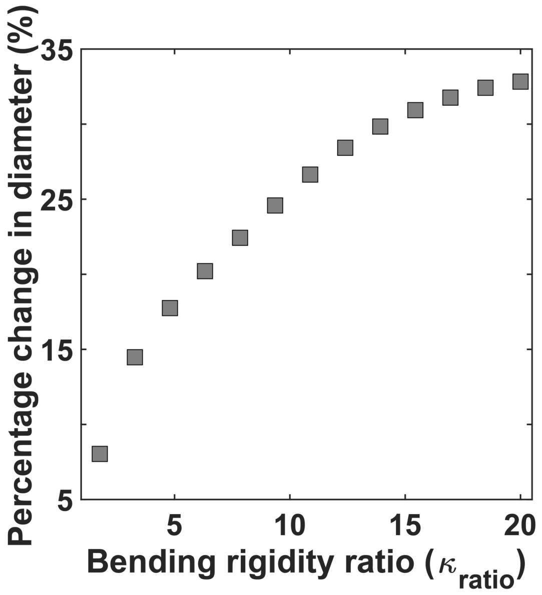


**Figure S4**: The tubule diameter increases with increasing the bending rigidity of the protein domain compared to the bare membrane. We defined the percentage of change in the tubule diameter as $(D - D_{\kappa_{ratio}=1})/D_{\kappa_{ratio}=1})$ and $\kappa_{\text{ratio}} = \kappa_{\text{proetin}}/\kappa_{\text{membrane}}$. Here, we set $C_0 = 3.5\ \mu m^{-1}$, $\lambda_0 = 0.9\ pN/\mu m$, $A_0 = 15\ \mu m^2$, and increased the bending rigidity ratio from $\kappa_{ratio} = 1$ (uniform rigidity) to $\kappa_{ratio} = 20$. Based on our results, with increasing the bending rigidity of the protein domain to $\kappa_{\text{ratio}} = 20$, the tubule diameter increases about 35% compared to the uniform bending rigidity. This increase in the tubule diameter with increasing the bending rigidity of the protein domain is qualitatively consistent with the experimental results shown in Figure 4g.
